# Germline-targeting SOSIP trimer immunization elicits precursor CD4 binding-site targeting broadly neutralizing antibodies in infant macaques

**DOI:** 10.1101/2023.11.07.565306

**Authors:** Ashley N. Nelson, Xiaoying Shen, Sravani Vekatayogi, Shiyu Zhang, Gabriel Ozorowski, Maria Dennis, Leigh M. Sewall, Emma Milligan, Dominique Davis, Kaitlyn A. Cross, Yue Chen, Jelle van Schooten, Joshua Eudailey, John Isaac, Saad Memon, Carolyn Weinbaum, Sherry Stanfield-Oakley, Alliyah Byrd, Suni Chutkan, Stella Berendam, Kenneth Cronin, Anila Yasmeen, S. Munir Alam, Celia C. LaBranche, Kenneth Rogers, Lisa Shirreff, Albert Cupo, Ronald Derking, Francois Villinger, Per Johan Klasse, Guido Ferrari, Wilton B. Williams, Michael G. Hudgens, Andrew B. Ward, David C. Montefiori, Koen K.A. Van Rompay, Kevin Wiehe, John P. Moore, Rogier W. Sanders, Kristina De Paris, Sallie R. Permar

**Affiliations:** Department of Pediatrics, Weill Cornell Medicine; New York, NY, USA; Human Vaccine Institute, Duke University Medical Center; Durham, NC, USA; Department of Integrative Structural and Computational Biology, The Scripps Research Institute, La Jolla, CA, USA; Department of Microbiology and Immunology, School of Medicine, University of North Carolina at Chapel Hill; Chapel Hill, NC, USA; Gillings School of Public Health and Center for AIDS Research, University of North Carolina at Chapel Hill; Chapel Hill, NC, USA; Department of Medical Microbiology, Academic Medical Center; Amsterdam, Netherlands; Department of Microbiology and Immunology, Weill Cornell Medicine; New York, NY, USA; New Iberia Research Center, University of Louisiana at Lafayette, New Iberia, LA, USA; California National Primate Research Center, University of California; Davis, CA, USA; Amsterdam Institute for Infection and Immunity, Infectious Diseases, Amsterdam, the Netherlands

## Abstract

A vaccine that can achieve protective immunity prior to sexual debut is critical to prevent the estimated 410,000 new HIV infections that occur yearly in adolescents. As children living with HIV can make broadly neutralizing antibody (bnAb) responses in plasma at a faster rate than adults, early childhood is an opportune window for implementation of a multi-dose HIV immunization strategy to elicit protective immunity prior to adolescence. Therefore, the goal of our study was to assess the ability of a B cell lineage-designed HIV envelope SOSIP to induce bnAbs in early life. Infant rhesus macaques (RMs) received either BG505 SOSIP or the germline-targeting BG505 GT1.1 SOSIP (n=5/group) with the 3M-052-SE adjuvant at 0, 6, and 12 weeks of age. All infant RMs were then boosted with the BG505 SOSIP at weeks 26, 52 and 78, mimicking a pediatric immunization schedule of multiple vaccine boosts within the first two years of life. Both immunization strategies induced durable, high magnitude binding antibodies and plasma autologous virus neutralization that primarily targeted the CD4-binding site (CD4bs) or C3/465 epitope. Notably, three BG505 GT1.1-immunized infants exhibited a plasma HIV neutralization signature reflective of VRC01-like CD4bs bnAb precursor development and heterologous virus neutralization. Finally, infant RMs developed precursor bnAb responses at a similar frequency to that of adult RMs receiving a similar immunization strategy. Thus, a multi-dose immunization regimen with bnAb lineage designed SOSIPs is a promising strategy for inducing protective HIV bnAb responses in childhood prior to adolescence when sexual HIV exposure risk begins.

## INTRODUCTION

In 2022, approximately 1.65 million [1.18 million-2.19 million] adolescents between the ages of 10 and 19 were living with HIV worldwide, and It is estimated that they account for around 10% of new adult HIV infections (*1*). Moreover, adolescents living with HIV have the highest rates of attrition in treatment and care, compounding the risk for transmission and disease among this group (*2, 3*). Therefore, it is imperative that HIV prevention efforts, including vaccine development, focus on this vulnerable population.

A successful HIV vaccine will need to elicit broadly neutralizing antibodies (bnAbs) capable of preventing viral entry of diverse HIV-1 variants. These bnAbs target relatively conserved epitopes on the HIV-1 envelope glycoprotein (Env) which include the CD4 binding site (CD4bs), the V3-glycan super site, the V2-glycan epitope on the apex of the trimer, the membrane-proximal external region (MPER) on gp41, and the interface of the gp120 and gp41 subunit (*4*). The induction of bnAbs to protective levels has continued present a major challenge in HIV vaccine development, and a variety of approaches are being employed to improve the elicitation of neutralizing antibody responses by immunization. HIV-1 Env SOSIP trimer-based immunization strategies represent one avenue of active exploration. Recently, it was shown that immunization with HIV-1 Env native-like SOSIP trimers can elicit antibodies with tier 2 autologous virus neutralization capacity in small animal models and rhesus macaques (*5–9*). In some cases, heterologous tier 2 virus neutralization responses have also been elicited, although these responses tend to be inconsistent among animals (*6*).

Retrospectives studies of human pediatric cohorts have revealed that children living with HIV generally develop bnAbs at a faster rate than adults living with HIV (*10–12*), suggesting that it may be easier to induce these responses by vaccination in the setting of the maturing immune system. Moreover, due to the period of relatively low risk of HIV acquisition following weaning, early childhood represents an opportune window for implementation of a vaccine strategy to elicit protective immunity prior to adolescence, when sexual HIV exposure risk begins. Initiating an HIV immunization regimen in infancy or early childhood also has the added benefit of allowing ample time for the multiple immunizations that will be required for the induction of bnAbs (*13, 14*). Notably, a vaccine capable of inducing bnAbs in children prior to sexual debut would be an important tool in the prevention of adolescent HIV (*15*).

Here, we report on the immunogenicity of wild type and B cell-lineage designed BG505 SOSIP trimers in infant rhesus macaques. The design of this SOSIP trimer was based on the BG505 clade A virus isolated from a 6-week-old infant who eventually developed a bNAb response within ∼2 years of infection (*10*). While the BG505 SOSIP.664 trimer can induce tier 2 autologous virus neutralizing antibodies to protective titers in adult rhesus macaques (*7*), it does not engage germ-line forms of HIV bnAbs (*16*). Thus, the BG505 SOSIP.v4.1-GT1 trimer was designed to improve the ability of this trimer to engage a broad range of germline precursor VRC01-class (CD4bs) and V1V2 apex bnAbs (*17*). Utilizing a second iteration of this trimer design, termed BG505 SOSIP.v4.1-GT1.1, we aimed to determine whether a combined BG505 SOSIP.v4.1-GT1.1 prime/BG505 SOSIP.664 boost immunization regimen could elicit bnAb responses in infant RMs.

## RESULTS

### Assessment of vaccine-elicited BG505 Env-specific IgG binding responses

We evaluated the immunogenicity of the wild type, BG505 SOSIP.664, which we will refer to as BG505 SOSIP, and the B cell lineage-designed, BG505 SOSIP.v4.1-GT1.1, which we will refer to as BG505 GT1.1 throughout the manuscript. Both SOSIP Env trimer immunizations were administered with 3M-052 adjuvant in squalene emulsion (3M-052-SE), as we previously demonstrated 3M-052-SE as an optimal adjuvant for eliciting robust antibody responses in the early life immune setting (*18*). **Figure 1A** provides a summary of the two immunization regimens evaluated in this study, which included a three-dose priming regimen with the BG505 SOSIP (n = 5) or BG505 GT1.1 (n = 5) initiated in the first week of life and immunizations 6 and 12 weeks later, followed by 3 boosts with the BG505 SOSIP immunogen at weeks 26, 52, and 78. Infant RMs developed high magnitude plasma IgG binding responses against the autologous BG505 SOSIP **(Fig. 1B; blue)** or BG505 GT1.1 vaccine antigen **(Fig. 1B; green)** after two immunizations (median ED_50_ at wk8: 7,516 and 20,003 for BG505 SOSIP vs BG505 GT1.1, respectively). Binding responses were boosted following each immunization and were maintained through week 96 **(Fig. 1B)**, the median AUC responses were similar between the two groups, but with greater variability among the BG505 GT1.1-primed infant RMs **(Fig. 1C)**. As the BG505 GT1.1-primed infants were later boosted with the BG505 SOSIP, we also measured plasma BG505 SOSIP-specific IgG binding responses in this group **(Fig. S1A)**. BG505 SOSIP-specific IgG binding responses were detectable at week 14, prior to receiving the first BG505 SOSIP boosts suggesting the development of antibody responses targeting overlapping epitopes on both immunogens. As expected, BG505 SOSIP-specific binding IgG responses increased following each boost yet, were consistently lower in magnitude compared to the BG505 GT1.1-specific binding responses **(Fig. S1A).** Not surprisingly, BG505 SOSIP-specific plasma IgG binding responses were consistently higher in the BG505 SOSIP-immunized group compared with the BG505 GT1.1-primed infants **(Fig. S1B)**. Antibody binding avidity was also evaluated by surface plasmon resonance. The avidity scores of BG505 SOSIP-specific IgG antibodies were similar between the vaccine groups **(Fig. S1C and S2)**. Thus, both immunization strategies were able to induce durable, high magnitude HIV-1 Env-specific IgG binding responses in infant RMs.

**Fig. 1.**
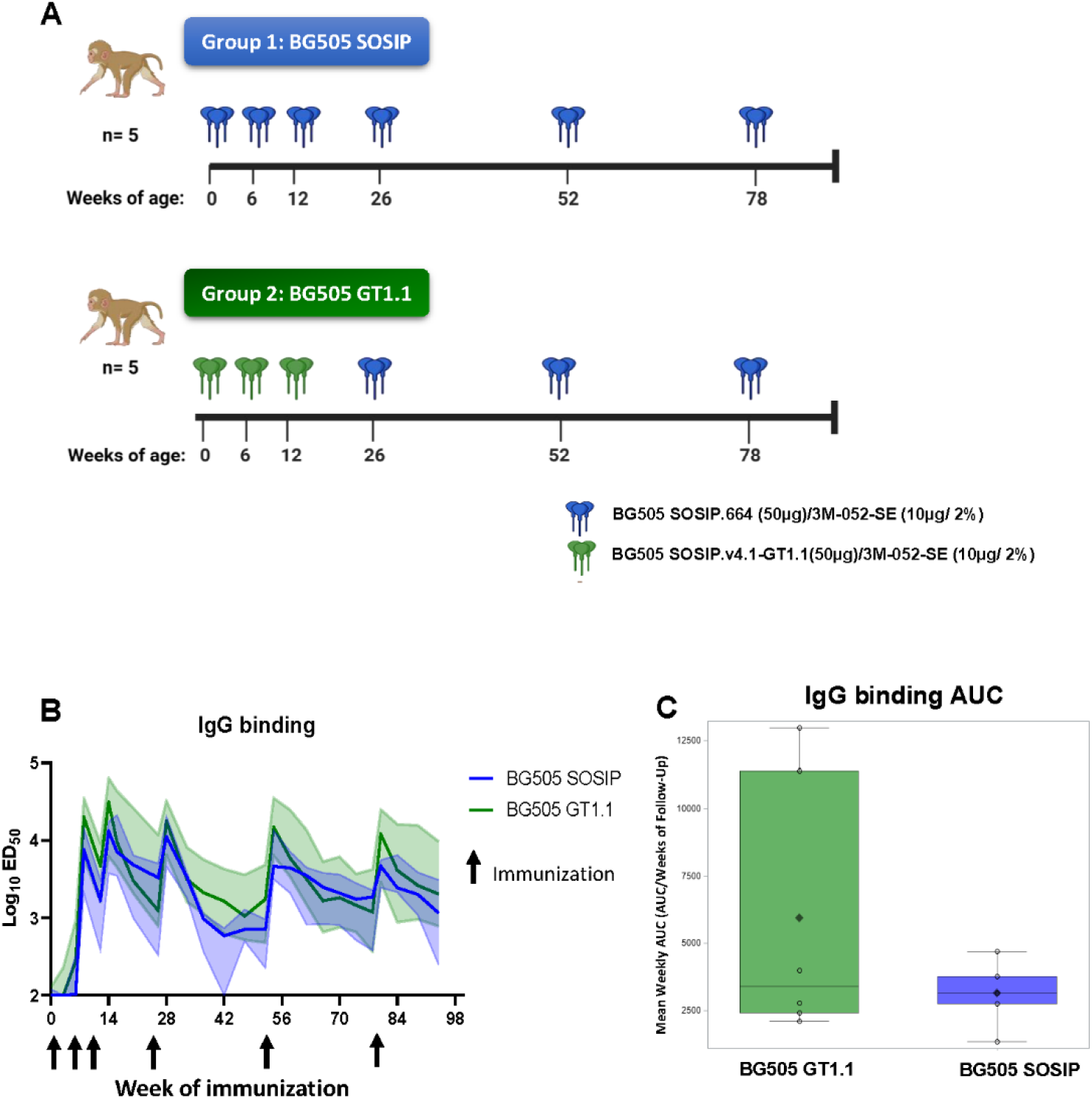
Development of durable, high magnitude Env-specific plasma IgG responses following BG505 SOSIP immunization. (A) Infant rhesus macaques (RMs) received 50μg of either BG505 SOSIP or germline-targeting BG505 GT1.1 (n=5/group) with the 3M-052-SE adjuvant. (B) Plasma IgG binding responses were measured to the matching immunogens BG505 SOSIP (group1; blue) or BG505 GT1.1 (group 2; green) for each group. Bold line represents the median, and shaded area represents the range. Black arrows indicate immunization time points. (C) Area under the curve analysis for vaccine-elicited plasma IgG binding responses. Individual datapoints are an empty circle. Group mean weekly AUC is a filled diamond. Q1/Median/Q3 make up the end and mid points of the box, with max and minimum values used to create the arms.

As a critical supporting response for B cell development, we next assessed the follicular T helper (Tfh) cell responses in lymph node mononuclear cells at week 54, 2 weeks after the 5^th^ vaccine dose. The proportion of antigen-specific Tfh cells were similar between the two groups (median BG505 SOSIP versus BG505 GT1.1: 1.51% and 1.35%, respectively**; Fig. S3A)**, consistent with observations of the similar magnitudes of the IgG binding responses. Further, we measured vaccine antigen-specific polyfunctional T cell responses via intracellular cytokine staining (ICS) in peripheral blood mononuclear cells collected after the 6^th^ dose of the vaccine regimen, at week 84, when adequate blood sample volume could be collected. Peripheral blood Env-specific CD4+ T cell responses were detectable at week 84, with a similar cytokine profile between the groups **(Fig. S3B)**. Env-specific CD8+ T cell responses were also elicited, with slightly higher production of IFNγ among the BG505 SOSIP-primed animals compared to the BG505 GT1.1-primed animals (median: 0.08% and 0.01%, respectively; **Fig. S3C)**.

### Neutralizing and non-neutralizing functions of BG505-specific antibodies

To assess the ability of the BG505 immunogens to induce autologous virus neutralizing antibodies (nAbs) in infant rhesus macaques, we screened sera against the autologous tier 1, BG505 GT1.1, and tier 2, BG505.T332N (*19*) pseudoviruses (PVs) (*20*). Tier 1 autologous virus neutralization responses were detectable in both groups after 3 immunizations (week 14; **Fig. 2A, B)**). As expected, median nAb titers against the BG505 GT1.1 virus were higher in the BG505 GT1.1-primed infants **(Fig. 2A)** compared to those immunized with BG505 SOSIP alone (median ID_50_ for BG505 GT1.1 vs BG505 SOSIP at week 14: 25,123 and 23, respectively; **Fig. 2A,B)**. In concordance, tier 2 autologous virus nAbs were detectable earlier in the BG505 SOSIP-only group, as 4 of 5 infants in the BG505 SOSIP-only group had detectable responses compared to 1 of 5 infants in BG505 GT1.1-primed group at week 14 **(Fig. 2C,D)**. After four immunizations (week 28), serum tier 2 autologous virus neutralizing antibodies were detectable in all animals from both groups and median responses peaked in both groups at week 80 (2 wks post 6^th^ immunization), corresponding with the boosted response in plasma IgG binding titers **(Fig. 1B)**. Overall, the BG505 immunogens were able to induce high titer, and durable autologous virus neutralization responses in infant RMs.

**Fig. 2.**
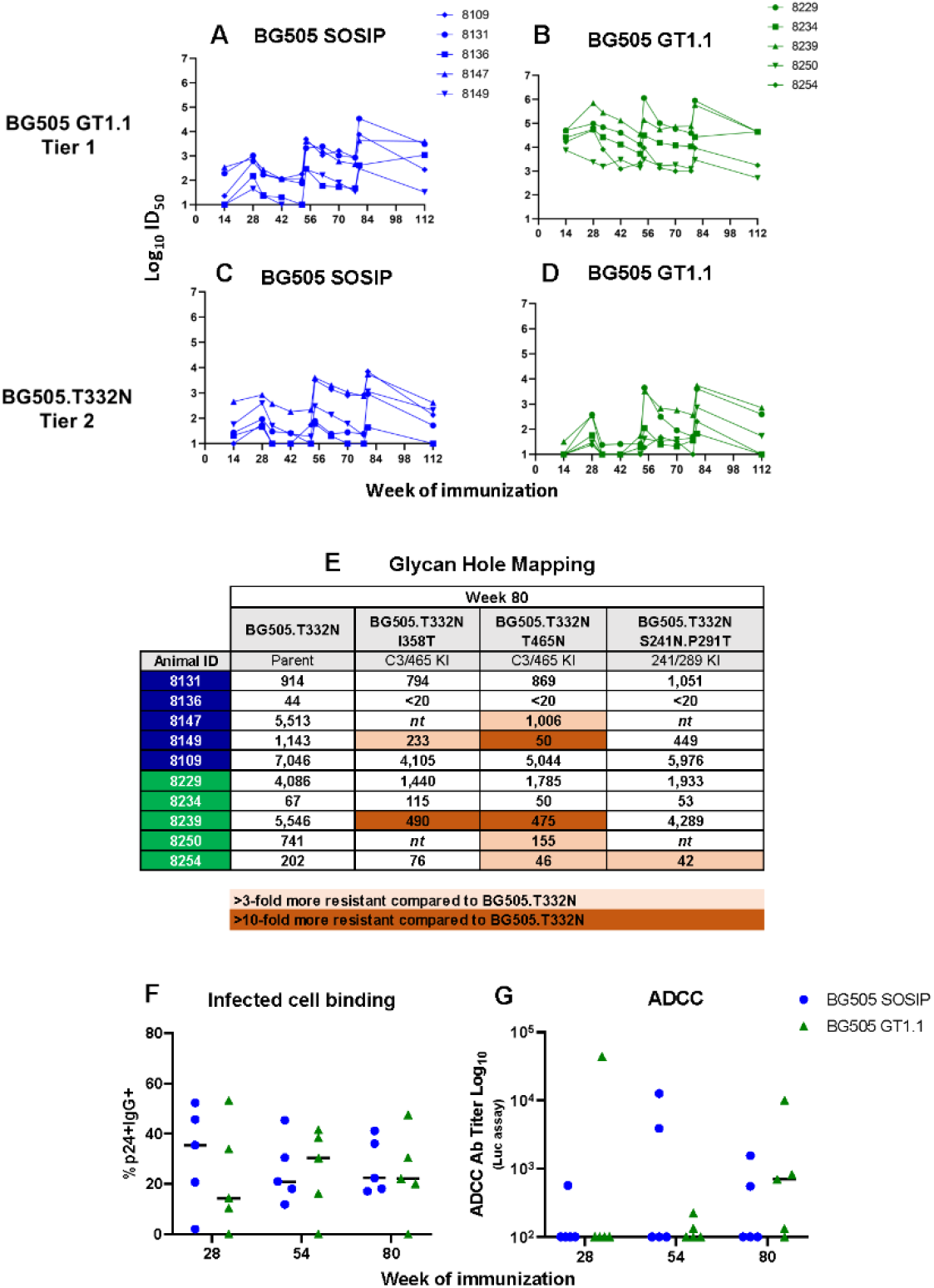
Autologous virus neutralization, recognition of infected cells, and ADCC mediating vaccine-elicited antibody responses in BG505 SOSIP-immunized infant RMs. (A-D) HIV-1 neutralization in sera was assayed against the BG505 GT1.1 tier 1 virus (A,B) or BG505.T332N tier 2 virus (C,D). (E) Sera at week 80 was assayed against a panel of mutant viruses to map neutralizing antibodies targeting glycan holes on the BG505 Env. Specificity was assigned based on a >3-fold reduction in ID_50_ to mutant virus compared to parent virus. (F) Levels of plasma infected cell IgG binding expressed as percent of BG505-infected cells with IgG binding. (G) Plasma ADCC activity against BG505 gp120-coated target cells. The plasma dilution endpoint antibody titers 2 weeks post the 4^th,^ 5^th,^ and 6^th^ immunizations for each animal are shown. In dot plots, horizontal lines represent median responses. For all graphs BG505 SOSIP immunized infants are in blue and BG505 GT1.1-immunized infants are in green.

In adult RMs immunized with BG505 SOSIP, tier 2 autologous virus nAbs generally target the C3/465 glycan hole on the BG505 envelope (*21, 22*). To map the specificity of the serum tier 2 autologous virus neutralization activity in infant RMs following immunization, we used a panel of mutant viruses on the parental BG505.T332N pseudovirus background. Each virus contained mutation(s) that filled one of the two major glycan holes identified on the BG505 Env at regions C3/465 (I358T or T465N), or 241/289 (S241N+P291T) (*23–25*). After 6 immunizations (week 80) 2 of 5 infant RMs receiving the BG505 SOSIP-only immunization strategy developed autologous tier 2 virus nAbs targeting the C3/465 glycan hole compared to 3 of 5 infants in the BG505 GT1.1-primed group **(Fig 2E).** One infant in the BG505 GT1.1-primed group also developed nAbs targeting the 241/289 glycan hole at week 80, although this did not persist through week 112 **(Fig S4)**, and 3 distinct infants exhibited this specificity of neutralizing antibody responses earlier at week 54 **(Fig. S4).** At 34 weeks after the last immunization (week 112), the C3/465 glycan hole remained the predominant target of autologous virus neutralizing antibodies **(Fig. S4)**.

While the SOSIP Env trimers are designed to elicit nAbs, we also evaluated other vaccine-elicited antibody functions, including Fc-mediated effector responses, which could contribute to protective HIV immunity in addition to neutralization. We detected the ability of vaccine-elicited IgG to recognize infected cells after the 4^th^ immunization in 4 out 5 animals from both immunization groups, with all RMs developing this response in the BG505 SOSIP-only group after the 5^th^ immunization (week 54; **Fig. 2F**). More specifically, among the BG505 SOSIP-only immunized group, the frequency of p24+IgG-bound cells was >20% in 3 out of 5 infants, whereas only 2 out of 5 GT1.1-immunized infants developed IgG responses capable of the same level of recognition of infected cells (**Fig. 2F; Fig. S5A-B)**. Interestingly, median infected cell-binding IgG responses peaked after 4 immunizations (week 28) in the BG505 SOSIP only group, and after 5 immunizations in the BG505 GT1.1-primed infants (week 54) (**Fig. 2F**). Furthermore, we observed inconsistent ADCC-mediating antibody responses against BG505 gp120-coated or infected target cells in both immunization groups **(Fig. 2G; Fig. S5C-F)**. Only one infant from each group exhibited ADCC responses after the 4^th^ immunization **(Fig. 2G)**. The response rate did not significantly improve after the 5^th^ and 6^th^ immunizations, with only 2 (BG505 SOSIP only) and 3 (BG505 GT1.1) infants developing ADCC responses by week 80 (2 weeks post 6^th^ immunization; **Fig. 2G**). Overall, both BG505 SOSIP immunization strategies induced antibodies that mediate autologous virus neutralization and can recognize infected cells, yet there was little induction of Fc-mediated antibody effector functions.

### BG505 GT1.1 priming is important for elicitation of CD4 binding site (CD4bs)-targeting antibodies

We used a bead-based antibody binding multiplex assay (BAMA) to determine if plasma IgG binding antibodies elicited following immunization with BG505 SOSIP alone, or priming with BG505 GT1.1 were specific to the CD4bs on the HIV Env. Thus, IgG binding to the BG505 GT1.1 SOSIP and the CD4bs mutant BG505 GT1.1/N279A.D368R was measured. In both immunization groups, the IgG binding responses at week 54 were similar against the two antigens **(Fig. 3A)**. A similar pattern was observed in the IgG binding responses to the 426c.DM.RS gp120 **(Fig. 3B)**, a recombinant protein demonstrated to elicit VRC01-class antibodies (*26*), and its corresponding CD4bs knock-out mutant, 426c.DM.RS.KO gp120 (*27*). Overall IgG avidity was also measured against the aforementioned 426c gp120 proteins, with two BG505 GT1.1-primed infants exhibiting a significant decrease in IgG avidity score against the 426 mutant protein **(Fig. 3D)**. Thus, although our BAMA results did not detect significant differences in binding between wild-type and CD4bs mutant antigens, SPR analysis revealed that BG505 GT1.1 priming seemed to elicit antibodies with a greater avidity for the CD4bs.

**Fig. 3.**
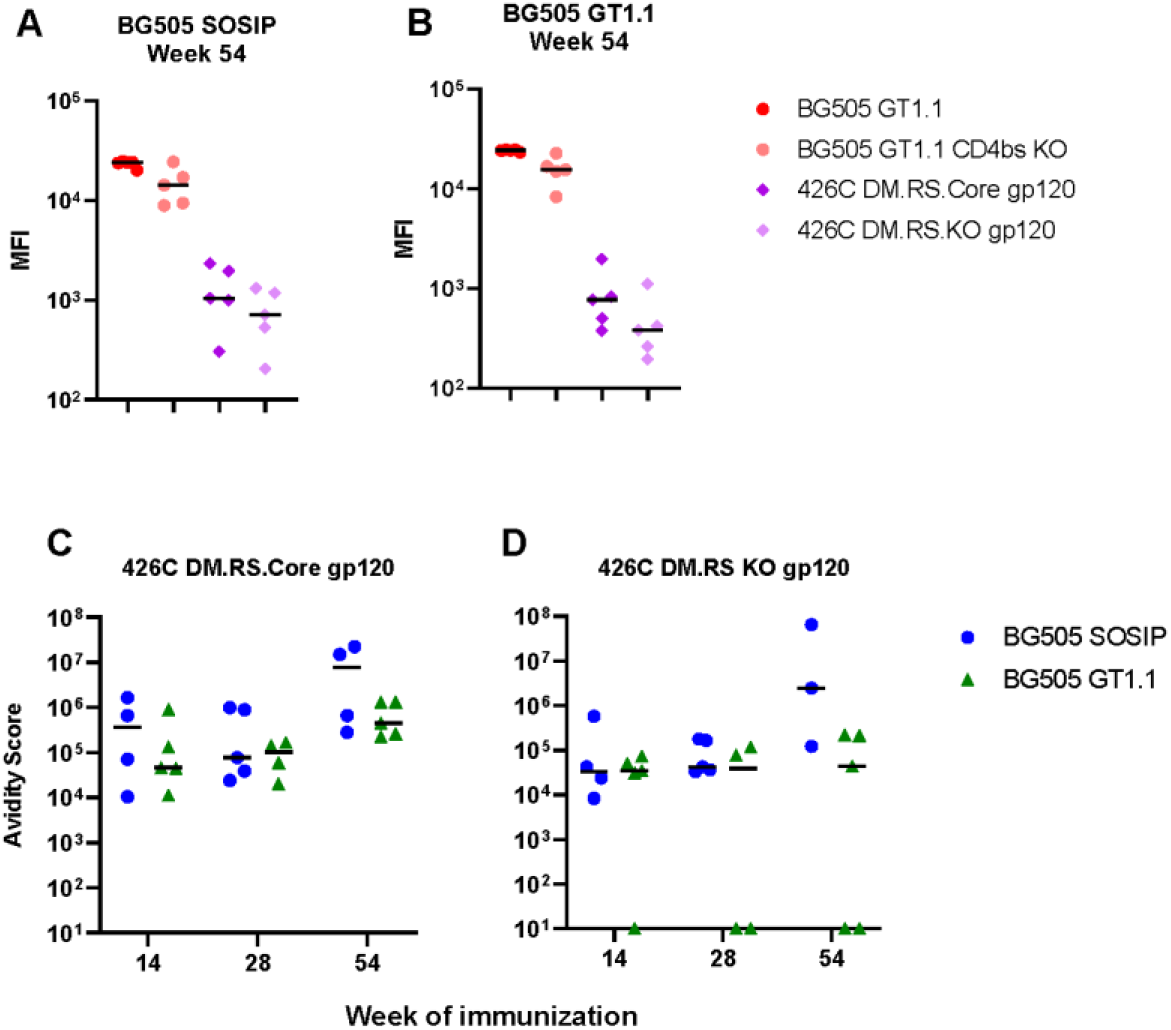
BG505 GT1.1 immunization elicits CD4bs-targeting binding antibodies in infant RMs. A multiplex bead-based assay was used to assess the specificity of vaccine-elicited plasma IgG binding for the CD4 binding site epitope on the HIV Env in BG505 SOSIP (A) and BG505 GT1.1 (B) immunized infant RMs. (C-D) Plasma IgG binding avidity scores as determined by SPR against the 426C DM.RS.core gp120 protein (C) and the corresponding CD4 binding site KO mutant (D).

To gain a more in-depth understanding of the epitope targets of the vaccine-elicited antibody responses, we utilized a negative-stain EM-based strategy to visualize polyclonal antibody binding (nsEMPEM) in plasma at week 80, two weeks after the final immunization **(Fig. 4)**. This nsEMPEM analysis revealed that circulating antibodies in animals from both immunization groups targeted various epitopes on the BG505 Env, with many that historically correlate with autologous virus neutralization. A total of five epitopes were common targets of antibodies among both groups: V1/V2/V3, C3/V5 region, gp120 glycan hole, the gp41 glycan hole and fusion peptide regions, and the trimer base **(Fig. 4A-B)**. One BG505 SOSIP-only immunized infant developed antibodies targeting the gp120-gp120 interface **(Fig 4A)**. Importantly, 4 out of 5 BG505 GT1.1-primed infants developed circulating antibodies targeting the CD4bs **(Fig 4B)**, with two infant RMs, 8229 and 8254, having autologous virus neutralizing antibodies that also mapped to the CD4bs **(Fig. S6**). Interestingly, BG505 GT1.1 primed infants generated antibodies that targeted a greater variety of epitopes at a higher frequency compared with BG505 SOSIP immunized infants **(Fig 4C-D)**. Overall, these data demonstrate that priming with the BG505 GT1.1 immunogen was essential for the elicitation of plasma CD4bs-targeting antibodies.

**Fig. 4.**
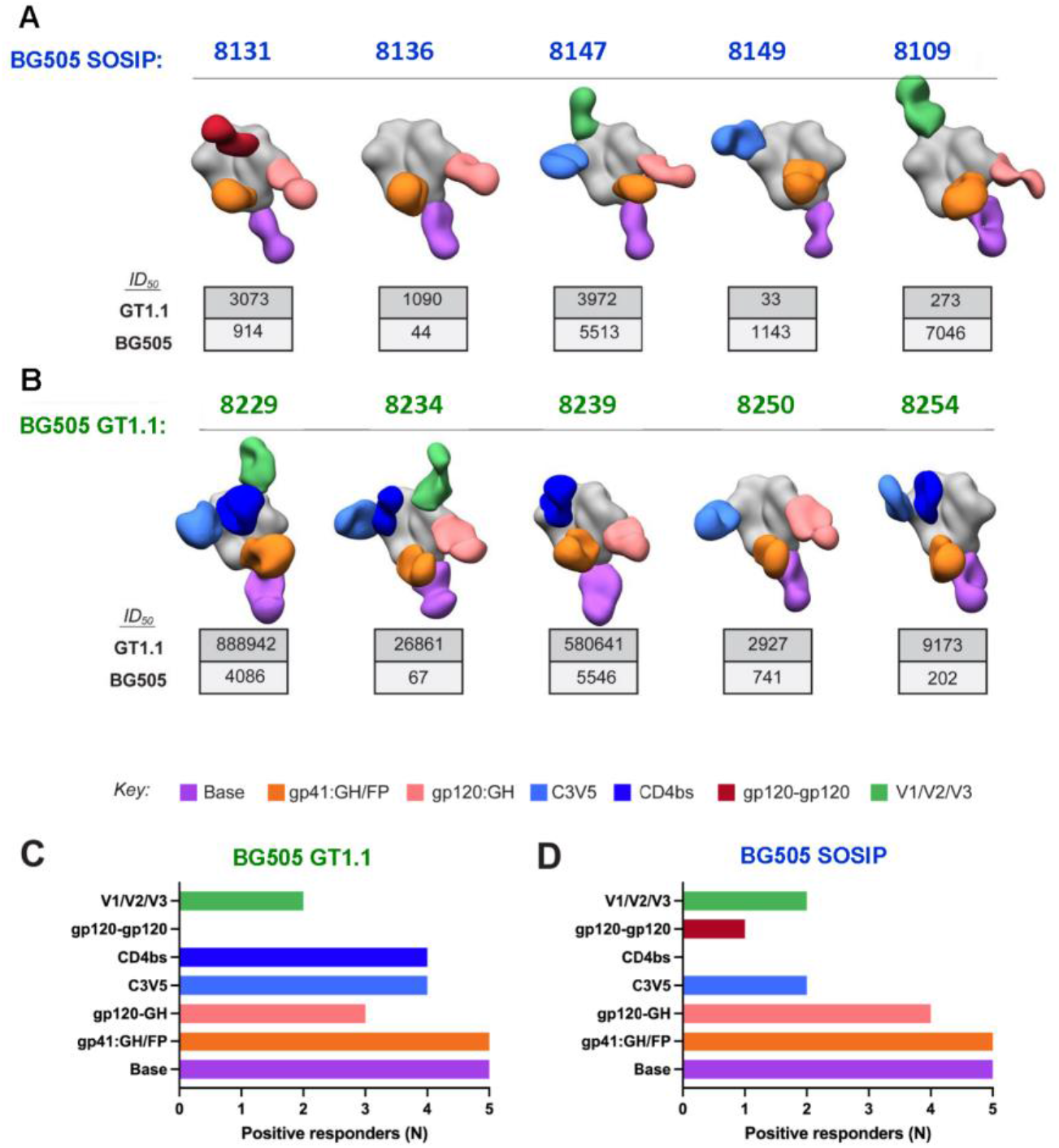
Visualization of polyclonal antibodies in plasma reveals CD4bs-targeting antibodies were elicited in the majority of BG505 GT1.1-primed infants. (A-B) nsEMPEM of either BG505 GT1.1 or BG505 SOSIP in complex with Fabs derived from plasma IgG at week 80. The neutralization ID_50_ titer against BG505 GT1.1 (tier 1 virus) and BG505.T332N (tier 2 virus) is shown in the table below each composite figure for each animal. Colors indicate previously defined epitopes on the BG505 SOSIP Env and are described in the key. (C-D) Summary of the proportion of epitope targets identified for each animal by nsEMPEM analysis for BG505 GT1.1 and BG505 SOSIP immunized infant RMs.

### BG505 GT1.1-priming induces the development of CD4bs-specific precursor bnAbs in infant RMs

The BG505 GT1.1 SOSIP was designed to engage VRC01-class bnAb precursors which target the CD4bs neutralizing epitope (*17*). Therefore, we assessed the development of CD4bs-specific bnAb precursor responses in the serum of the SOSIP-immunized infant RMs using a pre-defined neutralization phenotype indicative of VRC01-class bnAb precursors (*28*). Sera was screened for neutralization activity against the 426c.TM/293S/GnT1-virus, designed to engage germline-reverted forms of VRC01-class bnAbs, and the corresponding mutant virus containing the VRC01 resistance mutation D279K, 426c.TM.D279K/293S/GnT1-(*28*). A CD4bs bnAb precursor signature was assigned based on a 3-fold decrease in serum neutralization titers (ID_50_) against the mutant virus compared with the parental virus. After 4 immunizations, 2 of 5 BG505 GT1.1-primed infant RMs developed a neutralization signature indicative of plasma CD4bs-specific bnAb precursor development. This increased to a total of three infants by 6 immunizations **(Fig. 5A,B)**, and was maintained through week 112 **(Fig. 5C)**. In contrast, there was no detectable CD4bs bnAb precursor development observed among the BG505 SOSIP-only immunized infant RMs **(Fig. 5A,B)**. Thus, the B cell germline-targeting BG505 GT1.1 immunization strategy was superior to the wild-type BG505 SOSIP for the induction of CD4bs bnAb precursors, consistent with the EMPEM data.

**Fig. 5.**
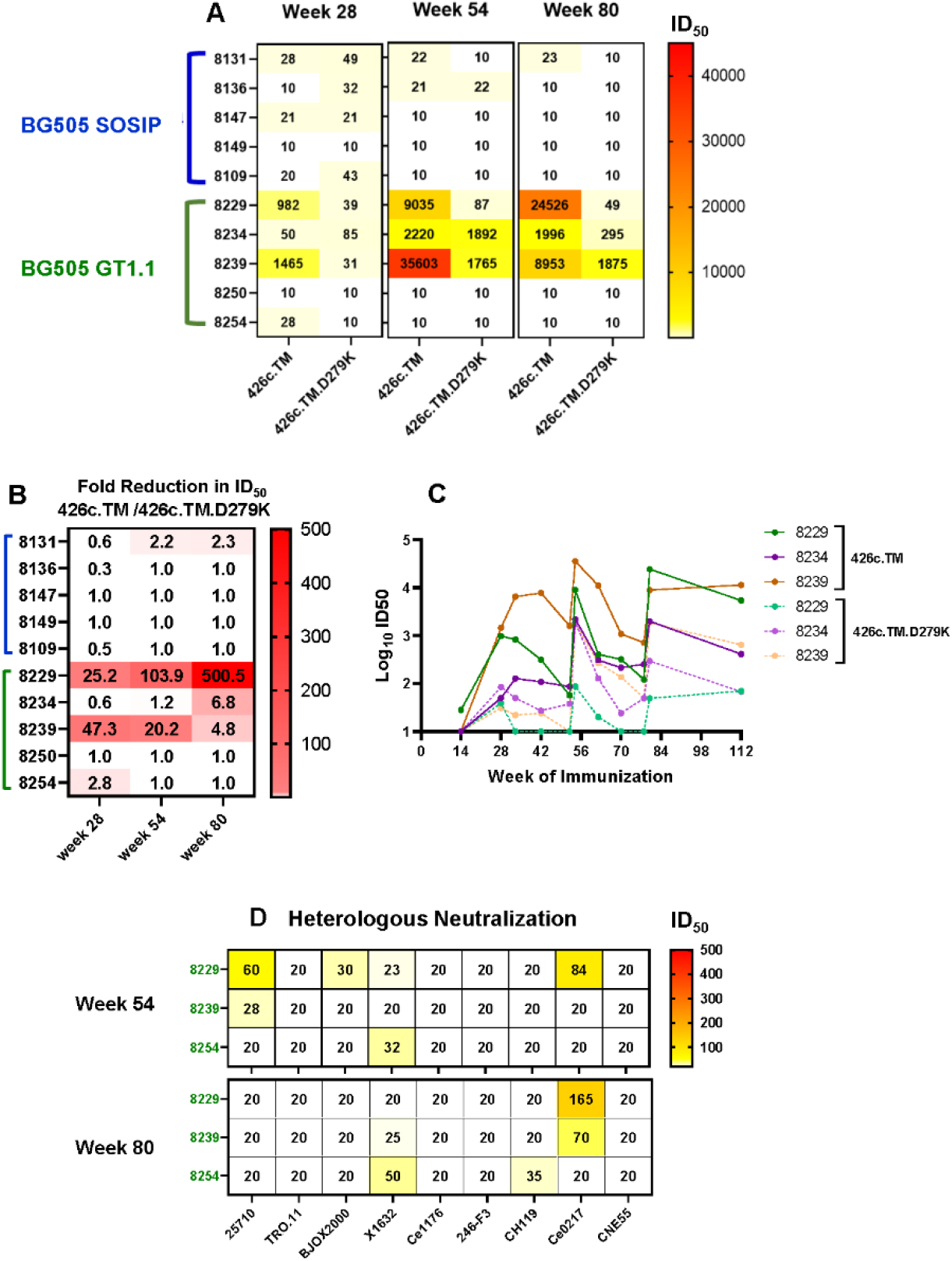
Three BG505 GT1.1-primed infants develop CD4bs-specific precursor bnAbs. Sera was screened for the development of VRC01-like CD4bs precursor bnAb responses using the 426c.TM/293S/GnT1-virus, which engages germline-reverted forms of VRC01-class bnAbs, and the CD4bs KO virus, 426c.TM.D279K/293S/GnT-. Neutralization ID_50_ titers (A) and fold decrease in ID_50_ against 426c parent and mutant viruses (B) are summarized in heatmaps from sera screened at 2 weeks post-4^th^, 5^th^, and 6^th^ immunizations. (C) Kinetics of the development of the CD4bs bnAb precursor signature in plasma of BG505 GT1.1 immunized infant RMs. (D) Three BG505 GT1.1-primed infants exhibited heterologous neutralization in plasma 2 weeks post-5^th^ and 6^th^ immunizations. Neutralization ID_50_ titers are summarized in the heatmap.

We also wanted to determine if any of the BG505 immunized infant RMs also developed heterologous virus neutralization responses. Plasma was screened at weeks 54 and 80 against a panel of tier 2 pseudoviruses representing global HIV-1 isolates (*29*). Heterologous virus neutralization responses were detectable in the plasma of 3 GT1.1-primed infant RMs (8229, 8239, and 8254), two of which also exhibited a plasma neutralization signature of CD4bs bnAb precursor development (8229 and 8239; **Fig 5D**). After 5 immunizations (week 54), animal 8229 exhibited cross-clade neutralization activity against 4 of the 9 viruses in the panel. The remaining 2 animals (8239 and 8254) neutralized one virus in the panel **(Fig. 5D)**. Across all animals, the ID50 titers were <100 against any viruses where neutralization activity was detectable. The breadth of neutralization responses in plasma did not improve after the final immunization, as by week 80 each animal exhibited neutralization activity against 1-2 viruses in the panel **(Fig. 5D)**. Interestingly, animal 8229 did exhibit an increase in neutralization potency against the Ce0217 pseudovirus **(Fig. 5D)**.

### Immunogenetics of the vaccine-elicited B cell repertoire of BG505 GT1.1-immunized infant RMs that developed a CD4bs bnAb precursor signature

To identify characteristics of the plasma neutralizing antibodies contributing to the development of CD4bs bnAb precursors in BG505 GT1.1-immunized infant RMs, we isolated and characterized mAbs from vaccine antigen-specific B cells of the three GT1.1-primed infants exhibiting this plasma bnAb precursor neutralization signature. Peripheral blood mononuclear cells (PBMC) cryopreserved at week 54 were single cell-sorted based on differential binding to the fluorescently labeled BG505 GT1.1 SOSIP and its corresponding CD4bs KO mutant (N279A.D368R) **(Fig. S7).** Because we were primarily interested in antigen-specific B cells with specificity to the CD4bs, B cells that bound to the BG505 GT1.1 SOSIP but not the CD4bs KO were collected. Among all recovered heavy (VH) and light (VL) chain transcripts, 26 functional pairs were identified **(Fig. 6A).** The heavy chain gene usage among these functional pairs was predominantly VH3 (n=12), followed by VH4 (n=6), and VH1 (n=7). Kappa chain gene usage was predominantly Vk1(n=6), while there was a greater variability in lambda (Vl) chain gen usage **(Fig. 6A)**. The median somatic hypermutation frequency among the VH genes was 9% **(Fig. 6B)**. The median CDRH3 length was 16 amino acids (AA), while the median CDRL3 AA lengths were 10 **(Fig. 6C-D)** and seemed to be within the range of those reported from other studies (*30, 31*).

**Fig. 6.**
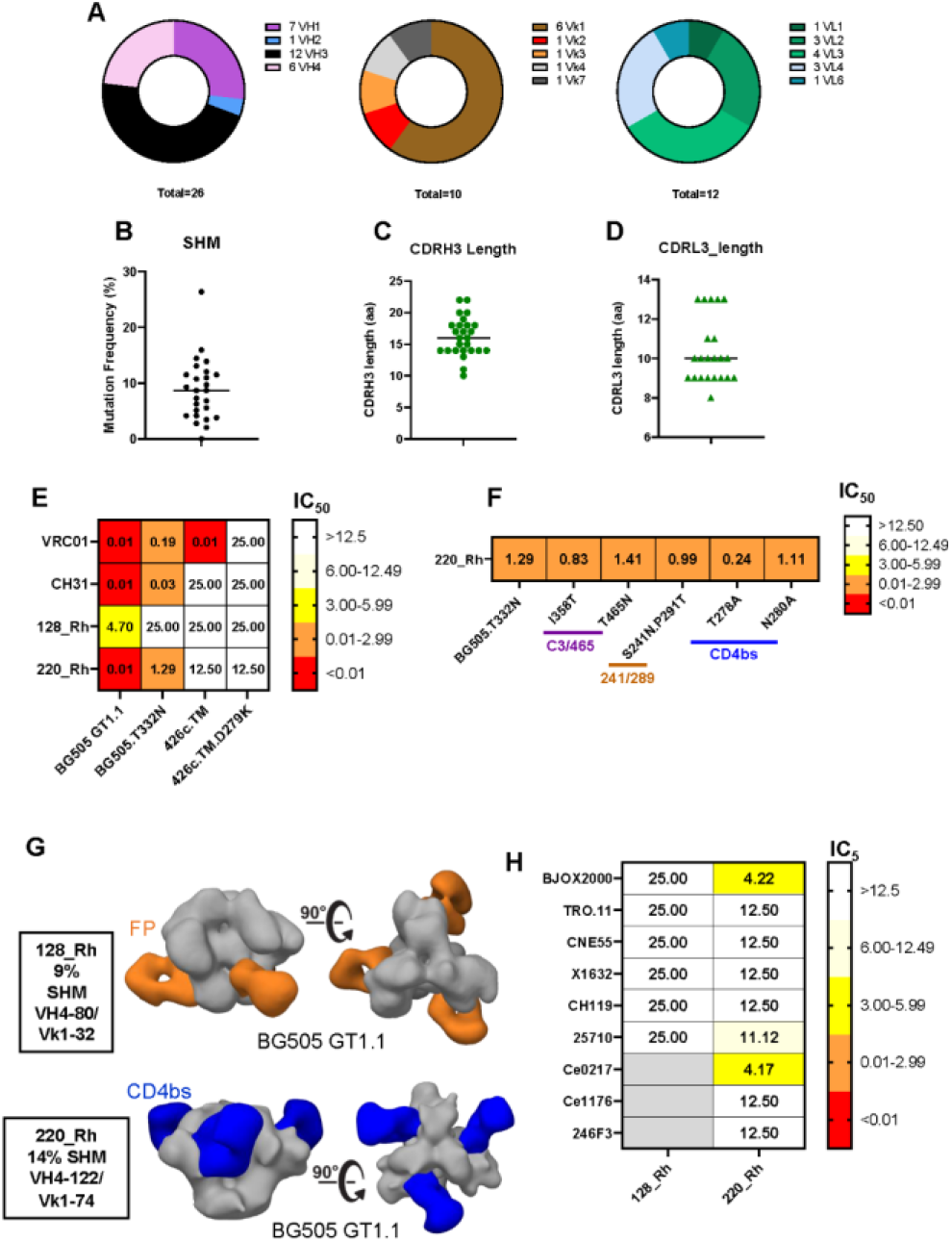
Characterization of mAbs isolated from vaccine antigen-specific B cells from BG505 GT1.1-primed infant RMs. BG505 GT1.1-specific B cells were single sorted from infants 8229, 8234, and 8239, and immunoglobulin heavy (Vh) and light chains, kappa (Vk) and lambda (Vl) regions were amplified and sequenced. Paired Vh and Vk/Vl sequences were obtained for 26 antibodies, and the proportion of each family is shown (A). Frequency of VH somatic hypermutation (B) and length of CDR3 among VH (C) and VL (D) genes of the isolated mAbs. (E) Heat map summarizing IC_50_ of two mAbs with autologous virus neutralization and CD4bs bnAb precursor neutralization capacity in comparison with HIV-1 bnAbs VRC01 and CH31. (F) Tier 2 autologous virus neutralizing mAb 220_Rh was assayed against a panel of mutant viruses to map epitope targets of neutralization activity. Specificity was assigned based on a >3-fold resistance to neutralization of the mutant virus compared to parent virus. (G) Negative stain electron microscopy epitope mapping of isolated rhesus mAbs to BG505 GT1.1 demonstrating binding to the fusion peptide region (orange) or the CD4bs (blue). (H) Heatmap summarizing neutralization IC_50_ against pseudoviruses of the global HIV neutralization panel.

For the 26 functional pairs identified, VH and VLs were cloned into expression plasmids and produced as recombinant mAbs. Each mAb was screened for binding to the BG505 SOSIPs. We were able to successfully generate 12 of the 26 mAbs identified. Of these 21 mAbs generated, 42%(n=9) bound to the BG505 SOSIP Env trimers **(Fig. S8)**. The mAbs positive for binding were then assessed for their ability to neutralize the autologous tier 1, BG505 GT1.1, and tier 2, BG505.T332N viruses. Two of the 14 mAbs screened exhibited potent tier 1 autologous virus neutralization ability, 128_Rh and 220_Rh **(Fig. 6E; Fig. S8C)**. Both mAbs originated from infant RM 8229 and were comprised of the VH4-80/Vk1-32 (128_Rh) and VH4-122/Vk1-74 (220_Rh) clonotypes. Four autologous virus neutralizing mAbs isolated from an adult RM immunized with the BG505 SOSIP were all comprised of the related VH4-39/Vk1.15 clonotype (*21*), suggesting a potential VH and VL gene usage signature for autologous virus neutralizing antibodies in RMs following immunization with BG505 SOSIPs. The isolated mAb 220_Rh, also exhibited potent tier 2 autologous virus neutralization capacity with an IC_50_ of 1.29 µg/ml **(Fig. 6E)** and was also more mutated in the heavy chain region compared to mAb 128_Rh (14% vs. 9%**; Table S3**). Moreover, mAb 220_Rh bound to the BG505 GT1.1 with greater affinity compared to 128_Rh as well as other non-neutralizing mAbs that were isolated from infant RMs (41_Rh and 113_Rh; **Fig. S9**). Interestingly, this mAb also bound to BG505 GT1.1 with more affinity than BG505 SOSIP, similar to VRC01. The stronger binding of VRC01 to BG505 GT1.1 than BG505 SOSIP is similar to what has been described for other BG505 SOSIP and germline adapted pairs of constructs (*14, 17*) and is due to the absence of glycans on the non-adapted trimer. When screened for the VCR01-like CD4bs bnAb precursor neutralization signature, neither mAb neutralized the 426c.TM/293S/GnT1-virus **(Fig. 6E)**.

The tier 2 autologous virus neutralizing mAb 220_Rh was also screened against mutant BG505.T332N pseudoviruses to determine the epitope specificity of its neutralizing activity. The neutralizing activity of this mAb did not target the C3/465 or 241/289 glycan holes **(Fig. 6F)**, despite the plasma neutralization activity of the corresponding RM having specificity for the C3/465 epitope at week 54 **(Fig. S4)**. However, a mutation in the CD4bs at position T278A resulted in increased sensitivity of the parent virus to neutralization by mAb 220_Rh **(Fig. 6F)**, consistent with the epitope mapping of neutralization activity in plasma of RM 8229 **(Fig. S6)**. Negative stain EM was performed to gain more insight into the epitope specificity the mAbs 128_Rh and 220_Rh **(Fig. 6G)**. Visualization of mAb 128_Rh complexed with BG505 GT1.1 demonstrated binding specificity for the gp41 fusion peptide region, while the epitope specificity of mAb 220_Rh mapped to the CD4bs **(Fig 6G)**. The two mAbs were also screened for heterologous virus neutralization activity. Importantly, mAb 220_Rh was able to neutralize 3 of 9 tier 2 viruses from the global neutralization panel (*29*) BJOX2000 (clade CRF07), Ce0217 (Clade C) and 25710_2.43 (clade C; **Fig. 6H**, analogous with the plasma heterologous neutralization activity of RM 8229 at week 54 **(Fig. 5E)**. Thus, mAb 220_Rh is an autologous virus neutralizing mAb with some heterologous virus neutralization breadth and specificity for the CD4bs initiated by germline-targeting SOSIP immunization in an infant RM.

### Comparison of neutralizing antibody responses elicited by BG505 SOSIP immunization in infant and adult rhesus macaques

Finally, we compared the vaccine-elicited neutralizing antibody responses of the infant BG505 SOSIP vaccinees to adult RMs that received similar BG505 SOSIP immunization strategies. The immunization schedules for the adult RM study are summarized in **Figure 7A**. Similar to the infants, one group of adult RMs received a BG505 SOSIP-only immunization strategy and the second group received a BG505 GT1.1 prime/BG505 SOSIP boost strategy **(Fig. 7A)**. Tier 1 and 2 autologous virus neutralizing antibody responses in plasma were compared at week 39 in the adult RMs to week 54 in the infant RMs, timepoints equivalent to 2 weeks post the 2^nd^ BG505 SOSIP boost among the BG505 GT1.1-primed cohorts. For the BG505 SOSIP-only immunized RMs, median responses against the autologous tier 1 BG505 GT1.1 virus was higher in the infants (median ID_50_: 2103) compared to adults (median ID_50_: 269) **(Fig. 7B)**. However, median responses were more similar across BG505 GT1.1-primed infant and adult RMs (median ID_50_: 31050 and 10677, respectively; **Fig. 7B**). The median tier 2 autologous virus neutralization responses against BG505.T332N were slightly higher in the infants (median ID_50_: 299) immunized with BG505 SOSIP only, compared with adults (median ID_50_: 76; **Fig 7C**), while the opposite trend was observed between the BG505 GT1.1-primed infants and adults (median ID_50_: 110 and 215, respectively; **Fig. 7C**).

**Fig. 7.**
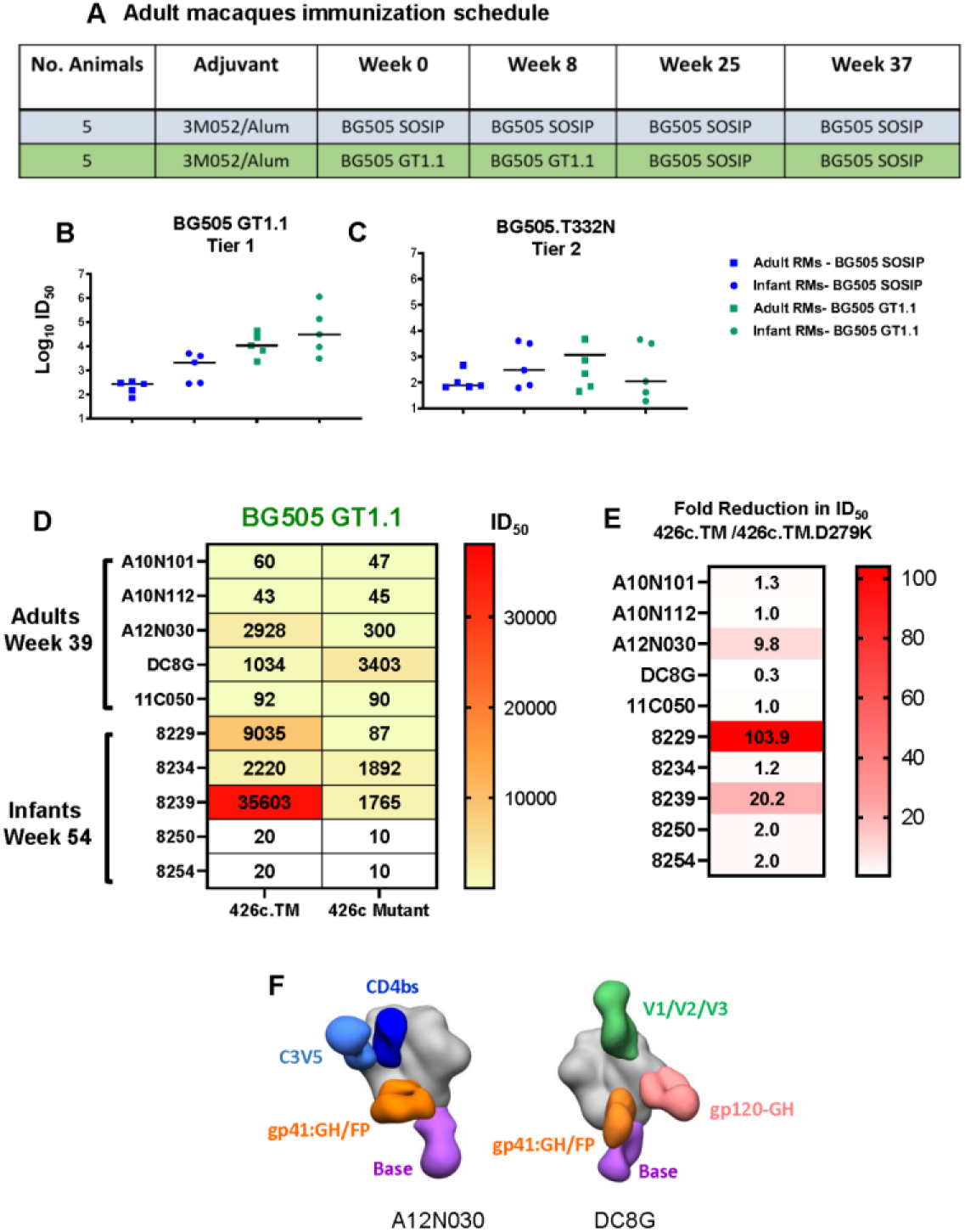
Comparison of neutralization responses elicited by BG505 SOSIP immunization in infant versus adult RMs. (A) Immunization schedule for adult rhesus macaques (RMs) who received either BG505 SOSIP or germline-targeting BG505 GT1.1 SOSIP trimer (n=5/group) with the 3M-052-Alum. Neutralization titers were compared between infant RMs at week 54 and adult RMs at week 39 against the BG505 GT1.1 (B) and BG505.T332N (C) PVs. In the dot plots, horizontal lines represent medians. (D) Sera was screened for the development of VRC01-like CD4bs-specific precursor bnAb responses among BG505 GT1.1 immunized infant and adult RMs. Neutralization ID_50_ titers are summarized in the heatmap for each animal. (E) Fold decrease in ID_50_ against 426c parent and mutant viruses are summarized in a heatmap. (F) nsEMPEM composite figure of two BG505 GT1.1-primed adult RMs with and without the CD4bs-specific bnAb precursor signature in plasma. Previously defined epitopes on the BG505 SOSIP Env are color coded.

Finally, we compared the frequency of CD4bs bnAb precursor development between the BG505 GT1.1-primed infant and adult RMs using the aforementioned 426c PV-based neutralization screening strategy **(Fig 5A)**. One out of the 5 adult RMs, A12N030, exhibited a neutralization signature indicative of CD4bs bnAb precursor development after 4 immunizations (week 39), compared with 2 of 5 infants, 8229 and 8239, that had developed this response (week 54, **Fig 7D,E)**. Analysis of polyclonal antibodies in plasma by nsEMPEM demonstrated differences in epitope targets of the vaccine-elicited antibodies between the one adult RM that developed the CD4bs precursor bnAb neutralization signature (A12N030) and a representative adult RM that did not (DC8G) develop this signature **(Fig 7F)**. Adult RM A12N030 developed antibodies targeting 4 different regions on the BG505 GT1.1 Env, including the CD4bs. Conversely, adult RM DC8G developed antibodies targeting 4 epitopes including two that were distinct from RM A12N030: V1/V2/V3, and gp120-GH. In contrast, infants that developed the CD4bs bnAb precursor signature in plasma responded to 4 or more neutralizing epitopes, including that of the CD4bs, after the 6^th^ immunization **(Fig. 4A and C)**. Together these data demonstrate infant RMs develop comparable neutralizing antibody responses to that of adult RMs receiving similar immunization regimens, yet potentially with more frequent epitope breadth.

## DISCUSSION

Immunization with BG505 SOSIP trimers have been successful at inducing high titer, tier 2 autologous virus neutralizing antibody responses in adult RMs (*5, 22*). Here we report on the immunogenicity of the BG505 SOSIP and germline targeting BG505 GT1.1 trimers in infant RMs. Immunization strategies incorporating either BG505 SOSIP or BG505 GT1.1 immunogens induced durable, high magnitude antibody binding and autologous virus neutralization responses, with little to moderate ADCC activity through 2 years. Moreover, we have shown that the germline targeting HIV Env trimer immunization strategy incorporating a BG505 GT1.1 prime/BG505 SOSIP boost regimen can induce precursor VRC01-like CD4bs bnAb plasma responses in infant RMs.

Previous work showed that human infant immune systems can respond to HIV antigens following immunization, with responses similar to or greater than adults (*32*). We have shown that autologous tier 2 virus neutralizing antibodies can be elicited in infant RMs by HIV Env trimer SOSIP immunization and reached levels comparable to that of adult RMs. Autologous virus neutralizing antibodies induced following BG505 SOSIP immunization have been shown to target holes on the BG505 env glycan shield at positions 241/289 (*5, 6, 24*) and within the gp120 C3-region (termed the C3/465 epitope) (*23*). Mapping studies revealed that infant serum autologous virus neutralizing antibodies targeted the 241/289 and C3/465 glycan holes early on, with C3/465 becoming dominant after 6 immunizations, consistent with observations in adult RMs (*21–23*). High tier 2 autologous virus neutralizing antibody titers have been associated with protection from SHIV challenge in adults, with a titer threshold of 1:500 (*5, 8*). Although we did not challenge infant RMs, we anticipate most would have been protected based on their neutralizing antibody characteristics, but further studies are needed to confirm this.

Most notably in this study, three BG505 GT1.1-primed infants exhibited a plasma HIV neutralization signature reflective of VRC01-like CD4bs bnAb precursor development. Negative stain electron microscopy-based polyclonal epitope mapping (nsEMPEM) of IgG in plasma further revealed that only infants primed BG505 5 GT1.1 developed plasma antibodies targeting the CD4bs (**Fig. 1**). Moreover, the frequency of development of this CD4bs bnAb precursor response in plasma as assessed by the 426c PsV based-neutralization screening strategy, was similar between BG505 GT1.1-primed infant and adult RMs after four immunizations. This was of particular interest because rhesus macaques do not have an ortholog for VH1-2, which is required to make VRC01-class bNAbs in humans (*33–35*). Together these data highlight the importance of germline-targeting immunogen priming in eliciting antibodies targeting the CD4bs, and potentially other bnAb epitopes, with the potential to develop into mature bnAbs. Further, one of the BG505 GT1.1-primed infants (8229) developed plasma neutralization activity against 4 of 9 global panel HIV-1 variants. As minimal heterologous virus neutralization activity was elicited in the other BG505 GT1.1-primed infants, future work will need to improve upon this immunization strategy to drive neutralization breadth development in more animals and advance early B cell lineages.

In characterizing the BG505 GT1.1 vaccine-elicited B cell repertoire in BG505 GT1.1-primed infants that developed a CD4bs bnAb precursor plasma neutralization signature, we isolated two mAbs with autologous virus neutralizing activity from the same BG505 GT1.1 SOSIP-primed infant (animal 8229) with the plasma heterologous virus neutralization breath. One of the mAbs, 220_Rh, exhibited epitope specificity for the CD4bs and also demonstrated some heterologous virus neutralization activity. Interestingly, this was one of the more somatically-mutated mAbs isolated from the vaccinated infants and had one of the longest heavy chain CDR3 lengths (Fig 6, Table S3). Moreover, this mAb did not exhibit a neutralization signature reflective of a VRC01-like CD4bs bnAb precursor based on screening against the 426c.TM PV panel, suggesting it may distinct from or evolved beyond the neutralization phenotype of VRC01-like bnAb precursor mAbs. Yet, this mAb demonstrates that mature neutralization lineages can be elicited by a germline-targeting HIV Env SOSIP in an early life immune system. As we did not isolate any mAbs with the VRC01-like CD4bs precursor bnAb signature despite detecting a strong plasma signature for this response, there may be a need to improve upon antigen selection for future B cell sorting strategies to identify these B cells. In all, defining the evolution pathway and how similar B cell lineages can be selected for via HIV immunization during the childhood immunization schedule should be the focus of future work seeking to initiate protective HIV immunity prior to adolescence.

This work demonstrates that immunizing RMs beginning in infancy with doses administered on a schedule that mirrors an under 5 childhood immunization schedule was able to elicit plasma neutralization and precursor bnAbs targeting the CD4bs at a similar frequency compared to immunized adult RMs. Further, B cell repertoire analysis revealed that mature nAbs can be elicited by germline-targeting SOSIP priming, following by wildtype SOSIP immunization. More importantly, these data highlight the potential for germline-targeting HIV immunization strategies beginning in infancy and the opportunity to shape these responses throughout early childhood to induce early bnAb lineages with the potential for development into mature plasma bnAb responses. Moreover, the results of this study are informative for designing clinical trials testing germline-targeting HIV immunization approaches in human infants.

## MATERIALS AND METHODS

### Ethics Statement

Type D retrovirus-, SIV- and STLV-1 free infant Indian origin rhesus macaques (RM) (Macaca mulatta) were maintained in the colony of California National Primate Research Center (CNPRC, Davis, CA). Animals were maintained in accordance with the American Association for Accreditation of Laboratory Animal Care standards and The Guide for the Care and Use of Laboratory Animals (*36*). All protocols were reviewed and approved by the University of California at Davis Institutional Animal Care and Use Committee (IACUC) prior to the initiation of the study.

### Reagents

The following reagents were obtained through the NIH HIV Reagent Program, Division of AIDS, NIAID, NIH: Peptide Pool, Human Immunodeficiency Virus Type 1 Subtype A (BG505) Env Protein, ARP-13122, contributed by DAIDS/NIAID; TZM-bl Cells, ARP-8129, contributed by Dr. John C. Kappes, Dr. Xiaoyun Wu and Tranzyme Inc. The 426C DM.RS.Core gp120, and 426C DM.RS.KO gp120 proteins were received from Leo Stamatatos (Fred Hutch Cancer Center).

### Infant rhesus macaque immunization and sample collection

Infant rhesus macaques (RMs) received 50µg of either BG505.SOSIP.664 (abbreviated to BG505 SOSIP) or the germline-targeting BG505 SOSIP.v4.1-GT1.1 (abbreviated to BG505 GT1.1) trimer (n=5/group) with 10µg 3M-052 adjuvant in a 2% squalene emulsion (3M-052-SE; provided by AAHI and 3M) at 0, 6, and 12 weeks of age. All 10 infant RMs were then boosted with the BG505 SOSIP at weeks 26, 52 and 78 of age. For sample collections, animals were sedated with ketamine HCl (Parke-Davis) injected at 10 mg/kg body weight. EDTA blood was collected via peripheral venipuncture. Plasma was separated from whole blood by centrifugation, and PBMCs were isolated by density gradient centrifugation using Ficoll®-Paque (Sigma).

### Adult rhesus macaque immunization schedule

Twelve healthy, uninfected (negative for simian immunodeficiency virus, simian retrovirus, simian T lymphotropic virus and Herpes B virus) male adult Indian origin rhesus macaques (Macaca mulatta) aged 7-12 years were used for these immunization studies (n=5/group). All animals were housed and maintained at the New Iberia Research Center (NIRC) of the University of Louisiana at Lafayette in accordance with the rules and regulations of the Committee on the Care and Use of Laboratory Animal Resources. The study protocol was reviewed and approved by the University of Louisiana at Lafayette Institutional Animal Care and Use Committee (Protocol #2016-8787-072). Animals were immunized with BG505 SOSIP (100 µg) or BG505 SOSIP GT1.1 (100 µg) at weeks 0 and 8, adjuvanted with TLR7/8 ligand 3M-052 (30 µg 3M-052/500 µg alum) and delivered subcutaneously. Animals were then boosted at weeks 25 and 37 with BG505 SOSIP (100 µg). All immunizations were delivered bilaterally as two doses of 50 µg.

### Surface Plasmon Resonance

IgG avidity was measured by surface plasmon resonance (SPR; BIAcore 3000, BIAcore/GE Healthcare) to a panel of HIV-1 antigens: BG505 SOSIP, BG505 GT1.1, 426C DM.RS.Core gp120 & 426C DM.RS.KO gp120. Binding responses were measured following immobilization by amine coupling of antigens (*37*) on CM5 sensor chips (BIAcore/GE Healthcare). Purified plasma IgG samples at 200 µg/ml were flowed (2.5 min) over spots (chip surfaces) of antigen followed by a dissociation phase (post-injection/buffer wash) of 10 min and a regeneration with Glycine pH2.0. Non-specific IgG binding of a pre-immune sample was subtracted from each post-immunization IgG sample binding data. Data analyses were performed with BIA-evaluation 4.1 software (BIAcore/GE Healthcare). Binding responses were measured by averaging post-injection response unit (RU) over a 10 s window; and dissociation rate constant, kd (second-1), was measured during the post-injection phase after stabilization of signal. Positive response was defined when both replicates have a RU value ≥10. Relative avidity binding score is calculated as follows: Avidity score (RU.s) = (Binding Response Units/kd) (*38*).

Binding of recombinant mAbs to BG505 SOSIP and BG505 GT1.1 were analyzed on BIAcore T200 instrument at 25 °C. Briefly, trimers were immobilized to RL values close to 50 response units (RU) by anti-His antibody that had been covalently coupled to a CM5 sensor chip as described (*39*). In each cycle, fresh Env protein was captured, and at the end of each cycle, Env trimer was removed by a pulse of 10 mM glycine (pH 2.0) for 90 sec at a flow rate of 50 μl min−1. Recombinant mAbs and bnAbs (VRC01 and PGT151) at a concentration of 0.5 µM was injected for 300 sec of association and 600 sec of dissociation, with a flow rate of 30 µl/min. Binding data were analyzed with BIAevaluation software.

### AIM assay

We performed the activation-induced marker (AIM) assay (*40*) on cryopreserved lymph node cells. Cells were thawed, washed, and rested in cRPMI at 37°C with 5% CO2 for three hours. Cells were then seeded into a 24-well plate at a density of 1.5-2×10^6^ cells per well before stimulation with BG505 Env peptide pool (1μg/mL), DMSO (Sigma-Aldrich), or SEB (0.5μg/mL) for 18 hours. Cells were washed, surface stained (Table S5) fixed with 1% PFA, and acquired within 4 hours. Data were acquired on an LSR Fortessa instrument running FACSDiva v8.0 software (BD Biosciences) and analyzed using FlowJo v10.8.1 (BD Biosciences).

### TZM-bl Neutralization Assays

Neutralization activity of serially diluted serum or recombinant mAbs (25 µg/ml or 12.5 µg/ml start) were measured in luciferase-expressing TZM-bl cells (NIH AIDS reagent) as previously described (*20*). Autologous virus neutralization responses were measured using the BG505 GT1.1 (tier 1) and BG505.T332N (tier 2) viruses. To map glycan hole and epitope specificity of neutralizing antibodies, a panel of viruses were used as summarized in **Figure S9**. Heterologous neutralization was assessed against a panel of 9 viruses from the global HIV neutralization panel (*29*). Samples were run in duplicate, and results were reported as the 50% inhibitory dilution or concentration (ID_50_ or IC_50_), which is the dilution of plasma or concentration of mAb resulting in 50% reduction in luminescence compared to that of virus control wells.

### Infected Cells Ab Binding Assay (ICABA)

The measurement of plasma Ab binding to HIV-1 envelope expressed on the surface of infected cells was conducted using flow-cytometry-based indirect surface staining according to methods similar to those previously described (*41*). Briefly, mock infected and the HIV-1 BG505 Infectious molecular clone (IMC)-infected CEM.NKR_ccr5_ cells were incubated with plasma samples at 1:100 dilution for 2h at 37°C, then stained with a viability dye (Live/Dead Aqua) to exclude dead cells from analysis. Subsequently, cells were washed, and permeabilized using BD Cytofix/Cytoperm solution. After an additional wash, cells were stained with FITC-conjugated goat-anti-Rhesus polyclonal antisera to detect binding of the plasma antibodies, and anti-HIV-1 p24 to identify infected cells. Cells positive for NHP plasma binding were defined as viable, p24 positive, and IgG-FITC positive. Results are reported as percent of p24+IgG+ positive cells and p24+IgG+ MFI among the viable p24 positive events after subtracting the background of the staining of the 2° antibody only and in addition to the subtraction of the background observed using the mock-infected cells and at baseline before immunization (week 0). We used the average of the baseline values from the other animals as baseline for the two animals whose baseline samples were not collected. Baseline correction for the FITC MFI was performed by subtracting pre-immunization MFI wherever applicable.

### ADCC

A flow-based (GTL) assay was used to test ADCC activity in plasma samples as previously described (*42*). Briefly, CEM.NKR_ccr5_ cells were coated with the recombinant BG505 gp120 in the GTL assay. Plasma samples were tested after a 4-fold serial dilution starting at 1:100. The cut-off for positivity in the GTL assay was >8% of Granzyme B activity. Plasma samples were also assessed using the Luciferase-based (Luc) ADCC assay against BG505 Infectious molecular clone (IMC)-infected CEM.NKR_ccr5_ cells. This is an ecto-IMC generated using the NL4-3 HIV backbone with the insertion of the HIV-1 BG505 envelope and the Luciferase reporter genes. The analysis of the results was conducted after subtracting the background detected with the pre-immunization samples. Similar to ICABA, we used the average of the baseline values from the other animals as baseline for the two animals whose baseline samples were not collected. After background subtraction, results were considered positive if the % specific killing was above 10%. ADCC endpoint titers were determined by interpolating the last positive dilution of plasma (>8% GzB activity).

### Antigen-specific B cell sorting

Biotinylated BG505 GT1.1 and BG505 GT1.1 N279A.D368R were coupled to streptavidin-conjugated AF647 (BioLegend) and VB515 (BioLegend), resulting in a fluorescently labeled probe. Cryopreserved PBMCs were washed and resuspended in PBS + 2% FBS. Cells were counted and incubated with labeled probes plus the antibody mixture summarized in **Table S2**. Sorts were carried out using a FACS Aria Cell Sorter and the following gating strategy: lymphocytes, singlets, live, CD14^-^, CD16^-^, CD3^-^, CD20^+^, IgD^-^, CD27^all^. We single-cell sorted IgD^-^ CD27^all^ memory B cells that were positive for binding to the BG505 GT1.1 but not the CD4bs KO antigen, BG505 GT1.1 N279A.D368R, into 96-well plates containing 12.5μl CDS sorting buffer as described in the SMART-Seq® HT Kit protocol (Takara Bio). Full-length cDNA amplification of single cells was performed per the kit protocol (Takara Bio). The analysis of the surface markers of the positive cells was performed on FlowJo.

### NGS and immunogenetics evaluation

Full length cDNAs were generated and amplified directly from a single cell through the SMART-Seq HT Kit (Takara Cat. No. 634437). Up to 200pg cDNAs were used to generate the dual index Illumina libraries using Nextera XT DNA Library Prep Kit (Illumina Cat No. FC-131-1096) and Nextera XT Index Kit v2 Set A (Illumina, Cat. No. FC-131-2001). Sequencing was performed on an Illumina NextSeq 500 sequencer to generate 2 × 76 paired end reads using NextSeq 500/550 High Output kit v2.5 (150 cycles) following the manufacturer’s protocol (Illumina, Cat. No. 20024912). The quality of cDNAs and Illumina libraries were assessed on a TapeStation 2200 with the high sensitivity D5000 ScreenTape (Agilent Cat, No. 5067-5592), and their quantity were determined by Qubit 3.0 fluorometer (Thermo Fisher). Reconstruction of paired heavy and light chains from short reads of B cell receptor (BCR) transcripts was performed using the software BALDR (*43*). As part of the BALDR pipeline, reads were mapped to the rhesus reference genome (MacaM v7) using STAR, and the four filtering strategies recommended in the BALDR documentation for enriching Ig and depleting non-Ig reads during rhesus BCR reconstruction were applied prior to assembly. The assembled sequences were then annotated using the IMGT database and IgBLAST, and subsequently mapped to the original reads with bowtie2. Redundant heavy and light chain sequences recovered by more than one filtering method were discarded and were reannotated using Cloanalyst software with the default Cloanalyst rhesus Ig library to determine B cell immunogenetics.

### Recombinant monoclonal antibody production

Antibody heavy chain and light chain genes were synthesized and cloned into plasmids containing the CMV promoter (GenScript). Final heavy and light chain plasmids were amplified, and paired segments were co-transfected in at a 1:1 ratio in Expi 293i Cells (Invitrogen) with transfection reagents (ExpiFectamine™ 293 Transfection Kit, ThermoFisher). After 4 to 5 days, cells were harvested, spun down, and supernatants were collected. Antibody was purified by Protein A Sepharose (GE Healthcare). Concentrations of purified mAbs was determined using a NanoDrop One (ThermoFisher).

### Negative stain EMPEM

Serum and sample preparation to obtain polyclonal fabs for electron microscopy were previously described (*44*). Briefly, IgG was isolated from 0.5 mL infant NHP sera of week 80 BG505 SOSIP (N=5) or BG505 GT1.1-primed group (N=5) using Protein G (Cytiva). Papain (Sigma Aldrich) was used to digest IgG to fabs. An overnight incubation using 15 µg of BG505 SOSIP or BG505 GT1.1 SOSIP trimer with 1 mg of fab mixture (containing Fc and residual papain) was performed and the complex was then purified the next day using a Superdex 200 Increase 10/300 GL gel filtration column (Cytiva). Purified complexes were concentrated and diluted to a final concentration of 0.03 mg/mL. The diluted samples were deposited on glow-discharged carbon coated copper mesh grids, followed by staining with 2% (w/v) uranyl formate. Electron microscopy images were collected on an FEI Tecnai Spirit T12 equipped with an FEI Eagle 4k x 4k CCD camera (120 keV, 2.06 Å/pixel), of an FEI Tecnai TF20 equipped with a Tietz F416 CMOS camera (200 kEv, 1.77 Å/pixel) and processed using Relion 3.0 (*45*) following the standard 2D and 3D classification procedures. UCSF Chimera (*46*) was used to generate the composite maps and the representative maps with identified epitopes have been deposited to the Electron Microscopy Data Bank

### Statistical Analysis

For comparison of IgG plasma binding data generated by ELISA, trapezoidal area under the curve (AUC) was calculated taking the average Y-value between each assessment point, minus the LLOD times the length of the period between assessments. Total AUC was calculated as the sum of all such trapezoids available in the data. The total AUC was divided by the number of weeks of follow-up available.

## Supporting information

All supplemental materials

## Acknowledgments

The authors would like to thank J. Watanabe and the staff of the CNPRC Colony Research Services for their support with these studies. Flow cytometry and cell sorting was performed in the Duke Human Vaccine Institute Flow Cytometry Facility (Durham, NC), and the UNC Flow Cytometry Core Facility. Thank you to Tom Bijl and Jonne Snitselaar of Amsterdam UMC, Academic Medical Center for the production and purification of the BG505 SOSIP trimers. We are also grateful for the PAVEG contract # HHSN272201800004C for supporting this work. The 3M-052-SE was kindly provided by AAHI and 3M. The funders had no role in study design, data collection and interpretation, or the decision to submit the work for publication. The content is solely the responsibility of the authors and does not necessarily represent the official views of the National Institutes of Health. Study data were collected and managed using REDCap (Research Electronic Data Capture) electronic data capture tools hosted at Duke University and Weill Cornell Medicine. Figure of study design was created with BioRender.com.

## Author contributions

S.R.P., K.D.P., K.K.A.V.R., K.W., M.G.H., J.P.M., R.W.S, D.C.M., R.D.,and A.N.N. contributed to conception and study design. A.N.N., X.S., S.V., G.O., M.D., S.Z., L.S., D.D. K.A.C., Y.C., J.V.S., K.R., J.E., J.I., S.M., A.B., S.C., S.B., K.C., A.Y., S.M.A., C.L., A.C., G.F., W.B.W participated in methodology, collection and/or analysis of the data. A.N.N., X.S., G.O., K.A.C., K.W., M.G.H., P.J.K., W.B.W., J.P.M., R.W.S., S.R.P., and K.D.P. contributed to the interpretation of the results. J.E., C.W., S.M.A., K.W., A.B.W., F.V., D.C.M., S.R.P., K.D.P., K.K.A.V.R. was involved in supervision or project administration. A.N.N. and S.R.P was responsible for drafting the original manuscript. All authors were involved in critical revision of the manuscript for accuracy and important intellectual content.

### Data and materials availability

All data are available in the main text or the supplementary materials. Any recombinant mAbs of interested may be made available with the completion of a material transfer agreement by contacting S.R.P. or A.N.N. Representative electron microscopy maps presented in this manuscript can be found in the Electron Microscopy Data Bank under accession codes EMD-41869, EMD-41870, EMD-41871, EMD-41872, EMD-41972, EMD-41973, EMD-41974, EMD-41975, EMD-41976, EMD-41977, and EMD-41978.

