## Supplementary material for "Germline-targeting SOSIP trimer immunization elicits precursor CD4 binding-site targeting broadly neutralizing antibodies in infant macaques": All supplemental materials

### Supplementary Materials and Methods

#### BG505 SOSIP Enzyme-linked immunosorbent assay (ELISA)

His-tagged BG505 SOSIP or BG505 GT1.1 SOSIP trimer was captured on nickel coated 96-well plates (Qiagen) at 0.15 µg/well and incubated overnight at 4℃. Plates were blocked with Blocking Buffer (2% Skim Milk in 1X TBS) and incubated for 1 hour at room temperature (RT). Plasma was serially diluted 3-fold starting at 1:100 in diluent (1X TBS, 20% Goat Serum, 2% Skim Milk) and incubated on the plate for 1 hour at RT. Control and recombinant monoclonal antibodies were run on each plate from 1000-0.5ng/ml. B12R1 (rhesusized CD4 binding site mAb) and PGT151 were used as positive controls, and 17b as a negative control. Plasma and recombinant mAbs were tested in duplicate. A polyclonal anti-monkey IgG horseradish peroxidase (HRP)-conjugate (Abcam) was used as the detection antibody and incubated for 1 hour at RT. KPL SureBlue Reserve system (VWR) was used for plate development and read at an absorbance of 450nm. GraphPad Prism was used to calculate ED50 (effective dilution at 50% response).

#### Binding Antibody Multiplex Assay (BAMA)

HIV-1 antigens were conjugated to polystyrene beads (Bio-Rad) as previously described (1) and incubated on filter plates (Millipore Sigma) for 30 minutes before plasma samples and controls were added. Plasma was diluted 1:500 in assay diluent (1% dry milk + 5% goat serum + 0.05% tween-20 in 1X phosphate buffer saline, pH 7.4.). Beads and diluted samples were incubated for 30 minutes, then IgG binding was detected using a PE-conjugated mouse anti-monkey IgG (Southern Biotech) at 4 µg/mL. Beads were washed and acquired on a Bio-Plex 200 instrument (Bio-Rad) and IgG binding was expressed as mean fluorescence intensity (MFI). To assess background, the MFI of binding to blank wells and non-specific binding of the samples to unconjugated blank beads were evaluated during assay analysis. An HIV-envelope specific antibody response was considered positive if above the lower limit of detection (100 MFI). To check for consistency between assays, the EC50 and maximum MFI values of the positive control was tracked by Levy-Jennings charts. The antigens conjugated to the polystyrene beads are as follows: BG505 SOSIP, BG505 GT1.1, BG505 GT1.1 N279A.D368R, 426C DM.RS.Core gp120 and 426C DM.RS.KO gp120.

**T cell phenotyping**

Cryopreserved PBMCs were thawed, washed, and resuspended in complete RPMI (cRPMI). Cells (1-2x$10^6$) were then stimulated for 6 hours with BG505 Env peptide pool (1µg/mL; NIH AIDS reagent), DMSO (Sigma-Aldrich), or Cell Stimulation Cocktail (0.5x, Invitrogen) at 37oC with 5% CO2. Brefeldin A (1x, Invitrogen) was added one hour into the incubation. After stimulation, cells were then permeabilized and stained with a panel of antibodies summarized in Table S4, and further described in (*36*).

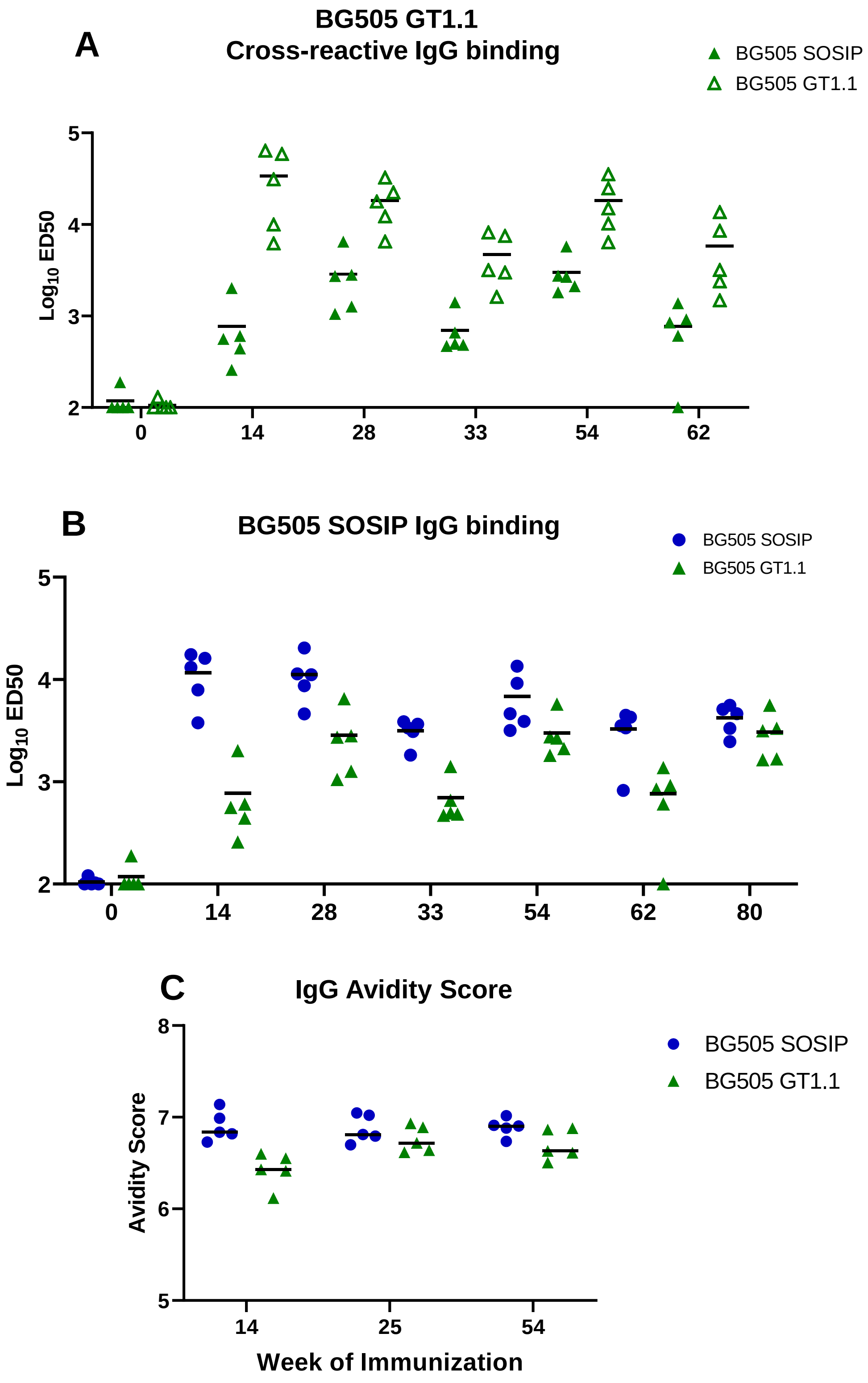


**Figure S1**. (A) Comparison of IgG binding to the BG505 SOSIP (closed triangles) and BG505 GT1.1 (open triangles) among  BG505 GT1.1-primed infant RMs. (B) Comparison of BG505 SOSIP-specific IgG binding responses among BG505 SOSIP (blue circles) and BG505 GT1.1 -primed infant RMs (green triangles). (C) Comparison of BG505 SOSIP-specific IgG binding avidity scores at weeks 14, 28, and 54 between the two immunization groups. In all dot plots, horizontal lines represent medians.

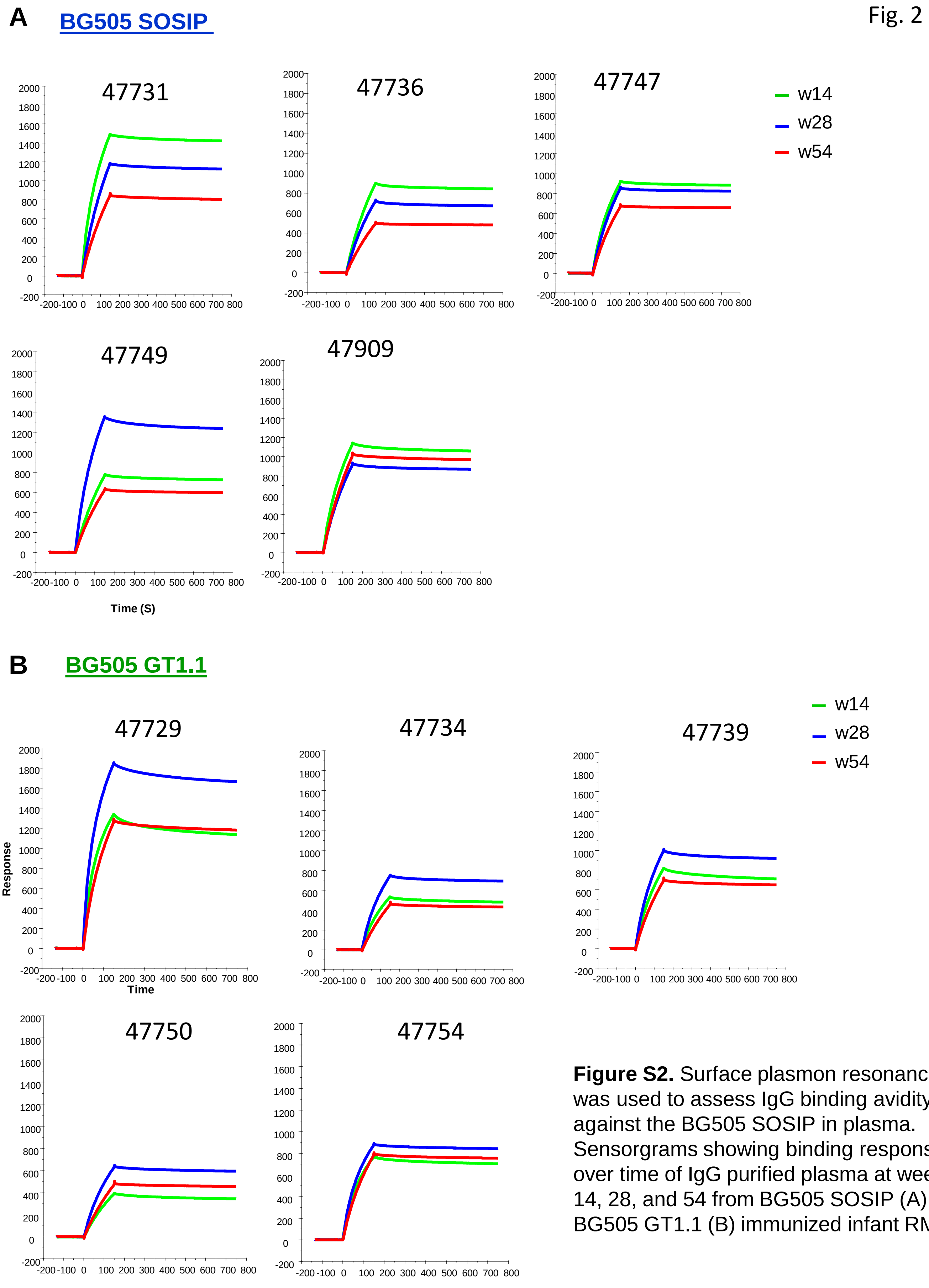


**Figure S2.** Surface plasmon resonance was used to assess IgG binding avidity against the BG505 SOSIP in plasma. Sensorgrams showing binding response over time of IgG purified plasma at weeks 14, 28, and 54 from BG505 SOSIP (A) and BG505 GT1.1 (B) immunized infant RMs.

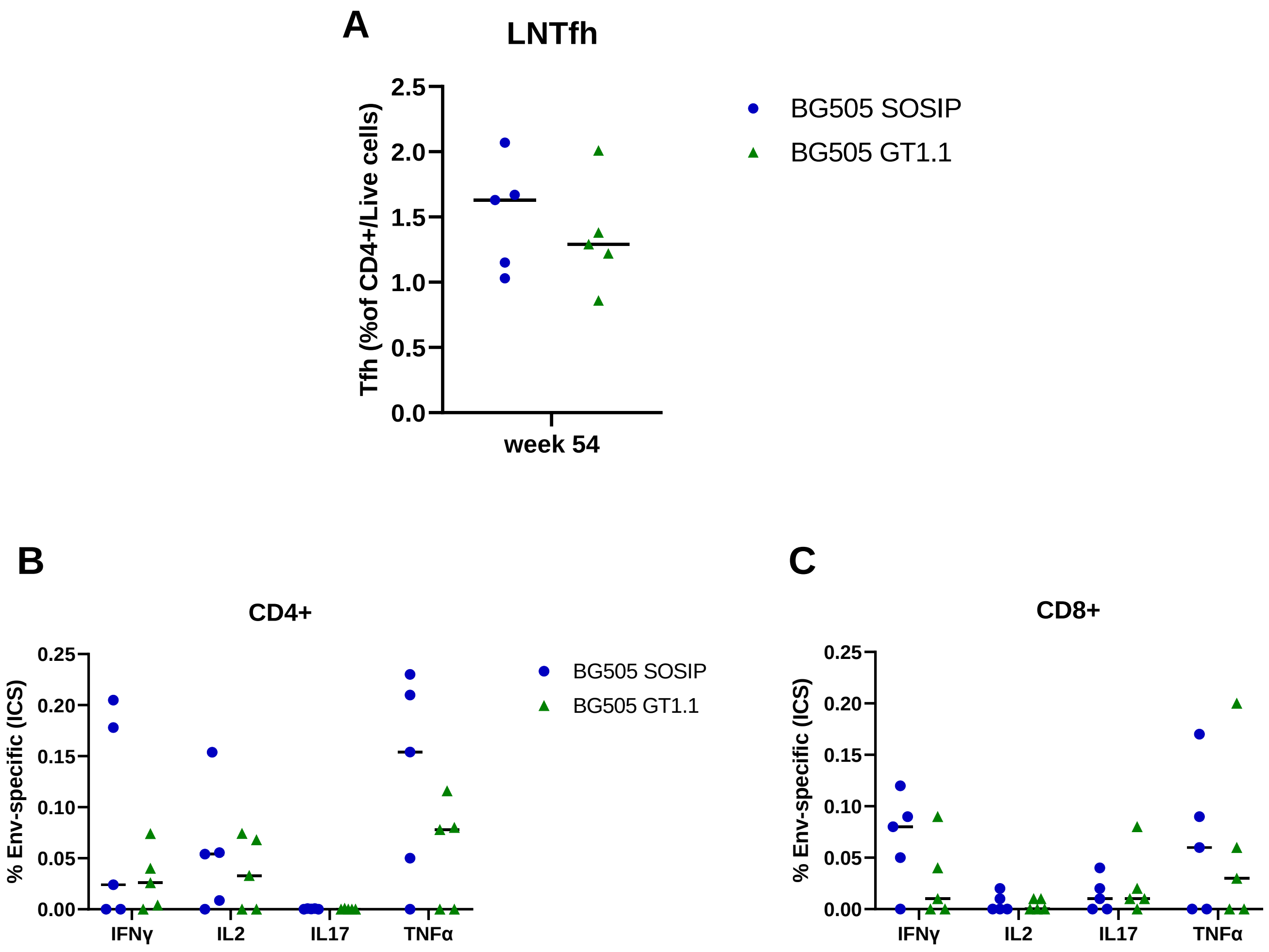


**Figure S3.** (A) BG505-specific Tfh responses were assessed in the lymph node (LN) after five immunizations at week 54. PBMC HIV-1 Env-specific CD4 (B) and CD8 (C) T cells were measured by ICS for IFNγ, TNFα, IL-17, and IL-2, five weeks following the 6th immunization.

| | wk54 | | | | wk112 | | | |
|---|---|---|---|---|---|---|---|---|
| | BG505/T332N | BG505/T332N I358T | BG505/T332N T465N | BG505/T332N S241N.P291T | BG505/T332N | BG505/T332N I358T | BG505/T332N T465N | BG505/T332N S241N.P291T |
| Animal ID | | C3/465 KI | C3/465 KI | 241/289 KI | | C3/465 KI | C3/465 KI | 241/289 KI |
| 8131 | 78 | 90 | *nt* | 39 | 52 | 134 | 62 | 55 |
| 8136 | 61 | <20 | 26 | <20 | <20 | <20 | <20 | <20 |
| 8147 | 3,475 | *nt* | *nt* | *nt* | 414 | 79 | 65 | 219 |
| 8149 | 299 | 108 | <20 | 92 | 210 | 86 | 27 | 219 |
| 8109 | 3,212 | 2,489 | 2,003 | 1,534 | 136 | 186 | 157 | 147 |
| 8229 | 2,579 | 462 | 507 | 985 | 398 | 164 | 168 | 146 |
| 8234 | 110 | 59 | 48 | 33 | <20 | 23 | 25 | 27 |
| 8239 | 2,865 | 61 | 64 | 3,544 | 720 | 81 | 65 | 907 |
| 8250 | 42 | <20 | <20 | <20 | 54 | <20 | 28 | 43 |
| 8254 | <20 | <20 | <20 | <20 | <20 | <20 | <20 | <20 |

>3-fold more resistant compared to BG505/T332N

>10-fold more resistant compared to BG505/T332N

**Figure S4.** Sera at weeks 54 and 112 were assayed against a panel of mutant viruses to map neutralizing antibodies targeting the glycan hole on the BG505 Env. Specificity was assigned based on a >3-fold reduction in $ID_{50}$ to mutant virus compared to parent virus. Table summarizes neutralizing $ID_{50}$ titers against each virus screened. Blue denotes infants immunized with BG505 SOSIP only, and green denotes infants primed with BG505 GT1.1.

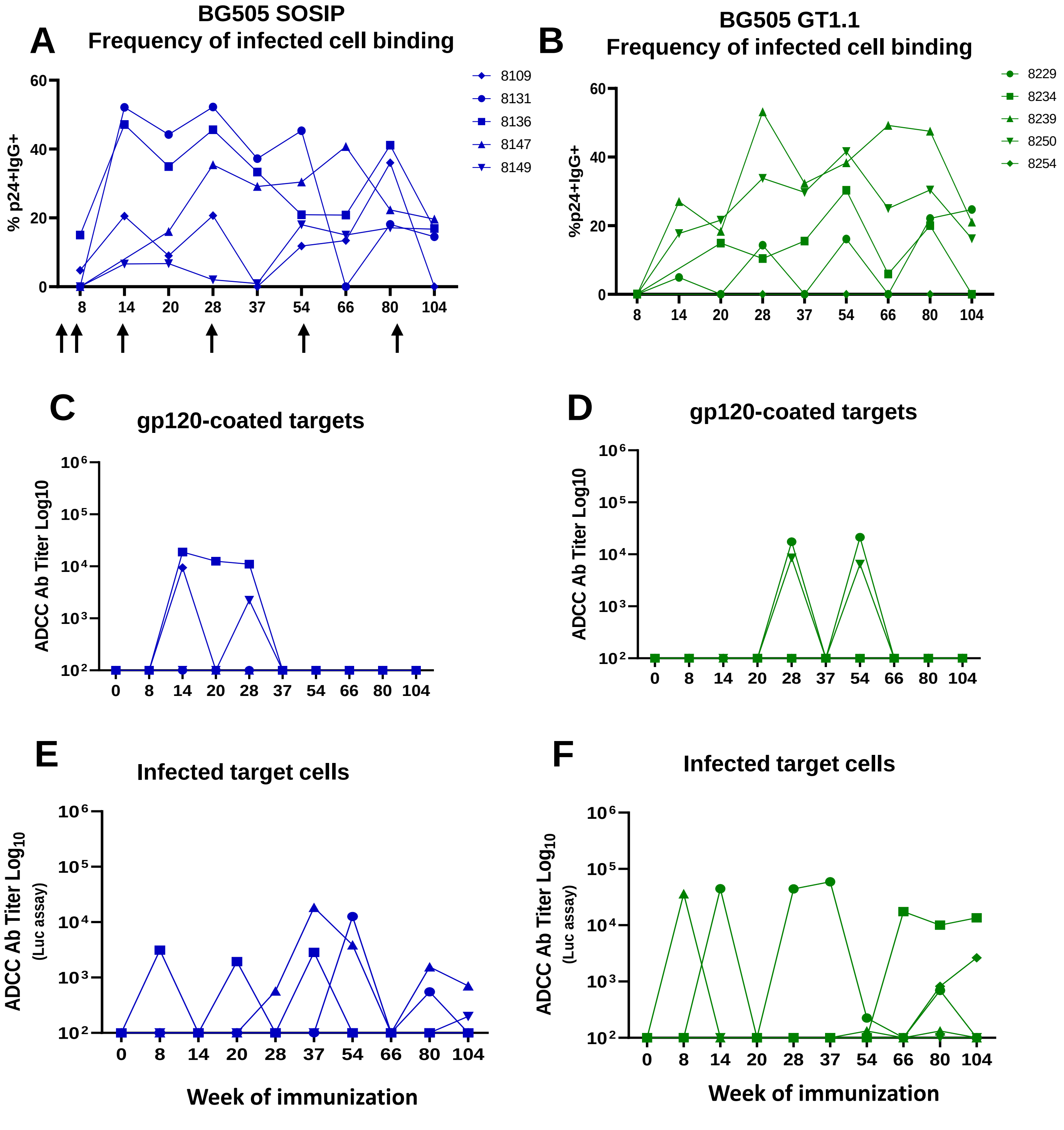


**Figure S5. ADCC antibody titers and infected cell binding antibodies.** Kinetics of endpoint titers of ADCC antibodies against BG505 gp120-coated target cells in BG505 SOSIP (A) and BG505 GT1.1 (B) immunized infants. ADCC endpoint antibody titers against BG505-infected cells in BG505 SOSIP (C) and BG505 GT1.1 (D) immunized infants. (E-F) The frequency of the infected cells recognized by the antibody responses, reported as %p24+IgG+ cells in immunized infants. Vaccination time points are indicated by the black arrows as referenced in panel A.

### Week 80

| | BG505.T332N | N160K | N301A | BG505 | T278A | N279A | N280A | BG505.G458Y | N295A | G354E | N611A |
|---|---|---|---|---|---|---|---|---|---|---|---|
| **Animal ID** | **Parent** | **V2 glycan** | **V3 glycan** | | **CD4bs** | | | | **2G12** | **C3/465** | **120-41** |
| 8131 | 807 | 1130 | 19508 | 904 | 2049 | 356 | 975 | 3740 | 1448 | 641 | 2122 |
| 8136 | *nt* | *nt* | *nt* | *nt* | *nt* | *nt* | *nt* | *nt* | *nt* | *nt* | *nt* |
| 8147 | 9408 | 6572 | 4046 | 4915 | 7396 | 9268 | 11161 | 10920 | 4345 | 687 | 7552 |
| 8149 | 1308 | 1052 | 1078 | 906 | 1404 | 1105 | 1266 | 25 | 1073 | <20 | 1334 |
| 8109 | 4356 | 5885 | 4063 | 6194 | 4429 | 4135 | 5330 | 12847 | 4681 | 6132 | 1989 |
| 8229 | 8215 | 6460 | 13836 | 6807 | >43740 | 9063 | 3810 | 659 | 6239 | 2572 | 6913 |
| 8234 | 11857 | 6199 | 8231 | 7620 | 21478 | 5245 | 8501 | 10174 | 6183 | 397 | 12172 |
| 8239 | 995 | 2040 | 897 | 878 | 1208 | 644 | 1128 | 831 | 784 | 49 | 1505 |
| 8250 | *nt* | *nt* | *nt* | *nt* | *nt* | *nt* | *nt* | *nt* | *nt* | *nt* | *nt* |
| 8254 | 157 | 215 | 391 | 139 | 248 | 169 | 38 | 2516 | 137 | 82 | 657 |

**>3 fold less sensitive compared to BG505.T332N**

*nt = not tested*

*all mutant viruses are on the BG505.T332N background unless otherwise noted

**Figure S6.** Sera at week 80 was assayed against a panel of mutant viruses to map neutralizing antibodies different epitopes on the BG505 Env. Specificity was assigned based on a >3-fold reduction in $ID_{50}$ to mutant virus compared to parent virus. Table summarizes neutralizing $ID_{50}$ titers against each virus screened. Blue denotes infants immunized with BG505 SOSIP only, and green denotes infants primed with BG505 GT1.1.

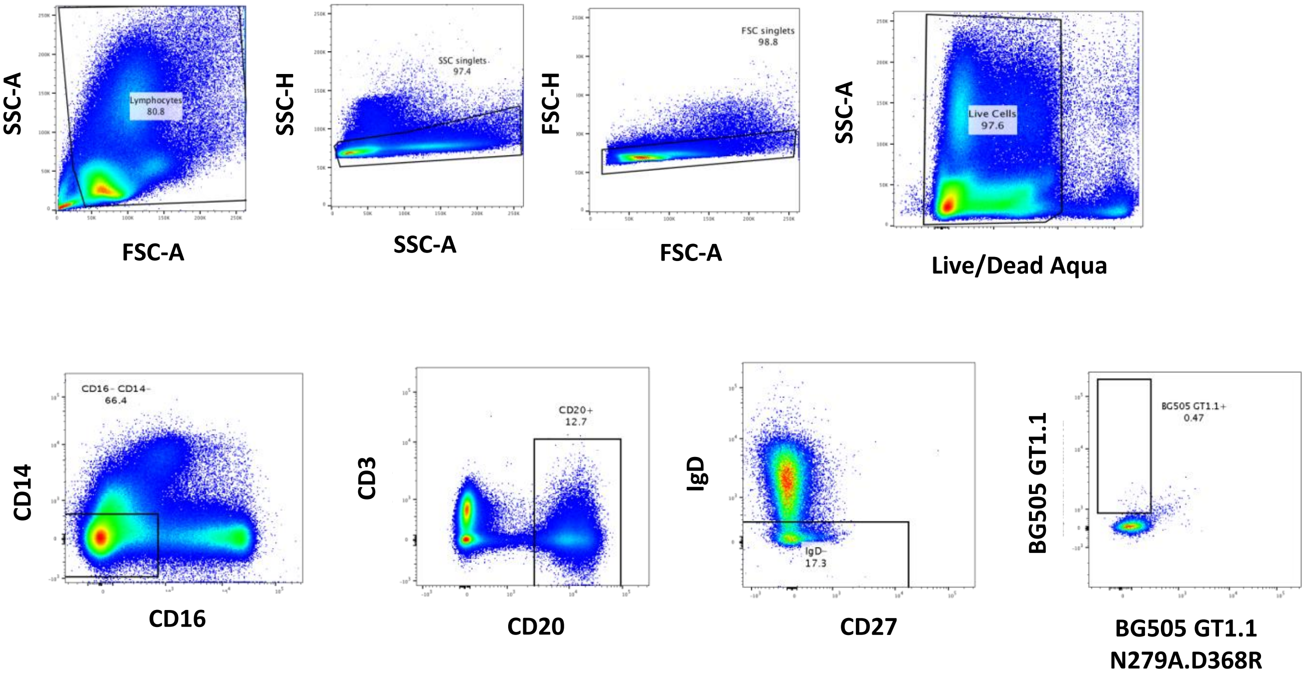


**Figure S7.** Representative flow plots showing the gating strategy for sorting BG505 GT1.1 specific B cells. Cells were first gated to identify lymphocytes, single, viable cells. Populations were further gated for CD14-, CD16-, CD20+ IgD- B cells. The final sort for antigen-specific B cells selected for differential binding to BG505 GT1.1 and the corresponding CD4bs mutant BG505 GT1.1 N279A.D368R. The final population of CD4bs targeting, BG505 GT1.1 specific B cells were single sorted into 96 well plates.

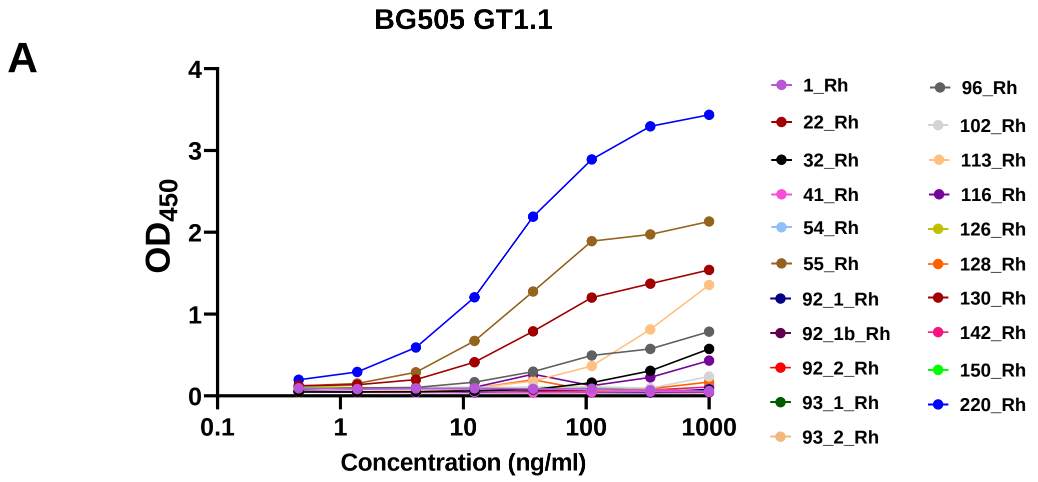


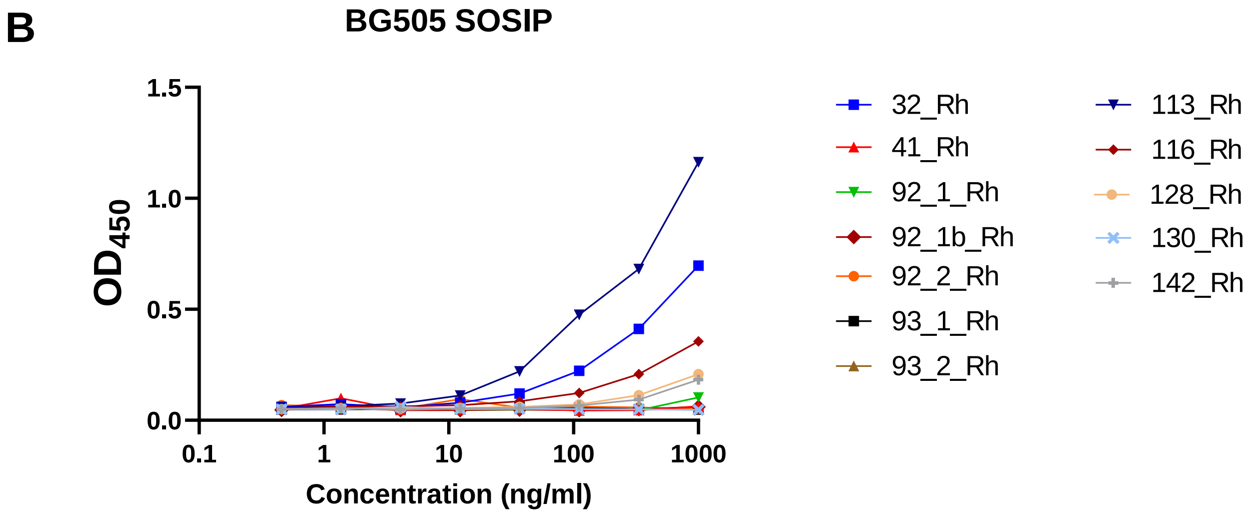


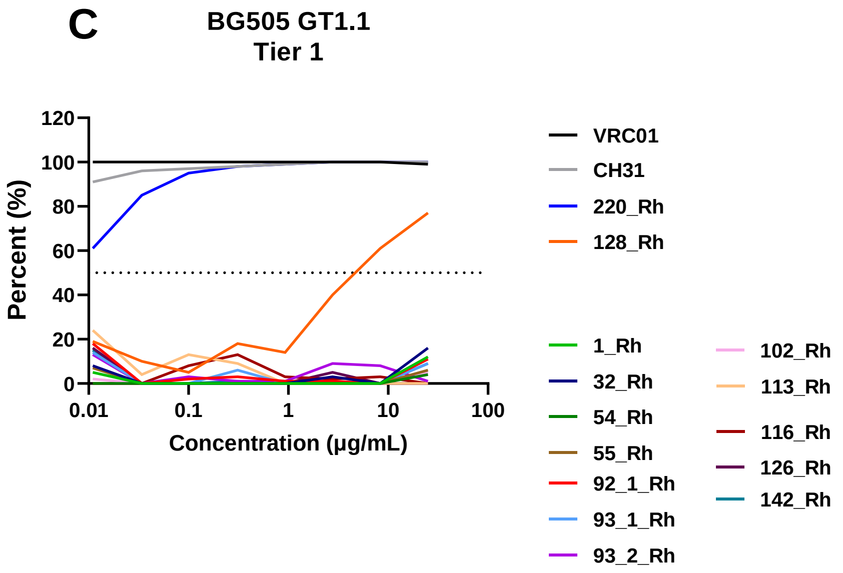


**Figure S8.** Graph shows the binding to BG505 GT1.1 (A) and BG505.664 SOSIP (B) of several mAbs isolated from infants. BG505 GT1.1 (tier 1) virus neutralization curves of selected mAbs (C).

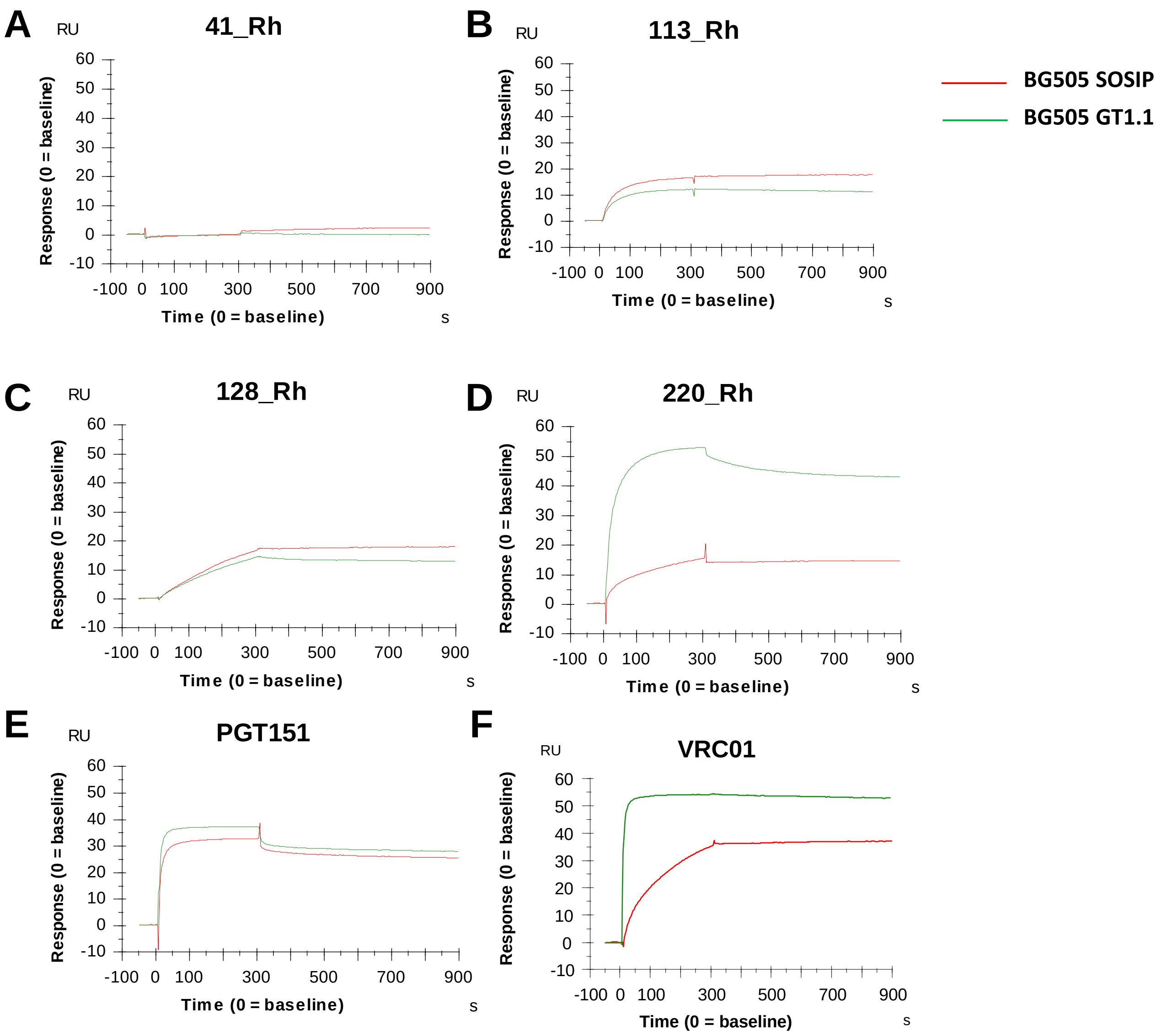


G

| | BG505 GT1.1 binding | BG505 GT1.1 Tier 1 Neutralization | BG505.T332N Tier 2 Neutralization |
|---|---|---|---|
| 41_Rh | - | - | - |
| 113_Rh | + | - | - |
| 128_Rh | + | + | - |
| 220_Rh | + | + | + |

**Figure S9.** Surface plasmon resonance was used to assess IgG binding avidity against BG505.664 and BG505 GT1.1 SOSIP of recombinant mAbs. Sensorgrams showing binding response over time of 4 isolated recombinant mAbs (A-D) in comparison to the HIV bnAbs PGT151 (E) and VRC01 (F). Summary of the ELISA binding and neutralization profiles of 4 mAbs screened for antibody avidity.

### A Glycan hole mapping panel

| Virus name | Epitope Target |
|---|---|
| BG505.T332N | Parent |
| BG505.T332N I358T | C3/465 glycan KI |
| BG505.T332N.T465N | C3/465 glycan KI |
| BG505.T332N.S241N.P291T | 241/289 glycan KI |

### B Epitope mapping panel

| Virus name | Epitope Target |
|---|---|
| BG505.T332N | Parent |
| BG505.T332N.N160K | V2 glycan |
| BG505.T332N.N301A | V3 glycan |
| BG505 | |
| BG505.T332N.T278A | CD4bs |
| BG505.T332N.N279A | |
| BG505.T332N.N280A | |
| BG505.G458Y | |
| BG505.T332N.N295A | 2G12 |
| BG505.T332N.G354E | C3/465 epitope |
| BG505.T332N.N611A | 120-41 |

**Figure S9.** Panel of HIV pseudoviruses used to evaluate glycan hole (A) and epitope specificity (B) of neutralizing antibodies.

**Table S1.** Summary of antigen-specific B cell sorting from three BG505 GT1.1-primed infant RMs that developed CD4bs bnAb precursor responses in plasma.

| Animal ID | Time point (week) | # total cells (input) | # of BG505 GT1.1-specific B cells recovered |
|---|---|---|---|
| 8229 | 54 | 9.5e6 | 240 |
| 8239 | 54 | 5.95e6 | 80 |
| 8234 | 54 | 3.8e6 | 160 |

**Table S2.** Panel of antibodies used for single B cell sorting

| Marker | Fluorochrome | Clone | Manufacturer |
|---|---|---|---|
| CD3 | PerCP Cy5.5 | SP34-2 | BD |
| IgD | PE | polyclonal | Southern Biotech |
| CD8 | PE Texas Red | 3B5 | Invitrogen |
| IgM | PE Cy5 | G20-127 | BD |
| CD16 | PE Cy7 | 3G8 | BD |
| Live / Dead | Aqua | N/A | Invitrogen |
| CD20 | BV650 | 2H7 | BioLegend |
| CD14 | BV570 | M5E2 | BioLegend |
| CD27 | APC Cy7 | O323 | BioLegend |

**Table S3.** Summary of immunogenetic characteristics of mAbs

| Animal ID | mAb ID | Hgene | CDRH3 length (aa) | %SHM | Lgene | CDRL3 length (aa) | %SHM |
|---|---|---|---|---|---|---|---|
| 8229 | 113_Rh | IGHV1-c*01 | 14 | 9 | IGLV3-l*01 | 13 | 10 |
| | 116_Rh | IGHV2-b*04 | 16 | 9 | IGKV1-b*01 | 9 | 6 |
| | **128_Rh** | **IGHV4-f*02** | **18** | **9** | **IGKV1-e*05** | **9** | **6** |
| | 130_Rh | IGHV3-e*01 | 22 | 7 | IGLV2-j*15 | 11 | 7 |
| | 142_Rh | IGHV4-f*03 | 16 | 12 | IGLV2-g*03 | 10 | 4 |
| | 150_Rh | IGHV1-a*03 | 18 | 11 | IGKV1-p*03 | 9 | 4 |
| | 171_Rh | IGHV4-n*01 | 20 | 13 | IGLV2-g*03 | 13 | 8 |
| | 32_Rh | IGHV1-c*01 | 14 | 10 | IGLV3-l*01 | 13 | 6 |
| | 41_Rh | IGHV3-v*02 | 15 | 11 | IGLV4-a*05 | 10 | 4 |
| | 60_Rh | IGHV1-h*02 | 18 | 11 | IGKV7-a*01 | 8 | 5 |
| | 92_Rh | IGHV3-y*02 | 14 | 4 | IGLV4-a*05 | 10 | 4 |
| | | IGHV3-j*02 | 14 | 2 | | | 0 |
| | | IGHV3-y*02 | 14 | 4 | | | 0 |
| | 93_Rh | IGHV3-y*02 | 10 | 6 | IGKV4-a*01 | 9 | 5 |
| | | IGHV3-y*01 | 22 | 4 | | | 0 |
| | | IGHV3-y*01 | 17 | 3 | | | 0 |
| | 96_Rh | IGHV4-m*02 | 13 | 5 | IGLV1-h*01 | 11 | 1 |
| | 199_Rh | IGHV4-j*02 | 17 | 3 | IGKV1-g*04 | 9 | 1 |
| | 202_Rh | IGHV3-r*03 | 11 | 6 | IGKV2-x*01 | 9 | 1 |
| | **220_Rh** | **IGHV4-n*01** | **19** | **14** | **IGKV1-p*01** | **9** | **7** |
| 8239 | 102_Rh | IGHV3-v*02 | 15 | 11 | IGKV1-p*03 | 9 | 5 |
| | 126_Rh | IGHV3-al*01 | 18 | 7 | IGKV3-f*07 | 10 | 3 |
| 8234 | 1_Rh | IGHV3-ai*01 | 17 | 0 | IGLV6-c*01 | 10 | 0 |
| | 22_Rh | IGHV1-h*02 | 14 | 14 | IGLV3-l*01 | 13 | 5 |
| | 54_Rh | IGHV1-h*03 | 20 | 26 | IGLV4-a*05 | 10 | 4 |
| | 55_Rh | IGHV1-e*03 | 14 | 16 | IGLV3-l*01 | 13 | 10 |

**Table S4.** Panel of antibodies used for T cell phenotyping

| Marker | Fluorochrome | Clone | Manufacturer |
|---|---|---|---|
| **Purified CD28** | N/A | L293 | BD |
| **Purified CD49d** | N/A | 9F10 | BD |
| **Viability** | Aqua | N/A | Invitrogen |
| **CD3** | APC-Cy7 | SP34-2 | BD |
| **CD4** | PE-CF594 | L200 | BD |
| **CD8** | BV786 | RPA-T8 | BD |
| **CD45RA** | V450 | 5H9 | BD |
| **CCR7** | PE-Cy7 | 3D12 | BD |
| **IL-2** | PerCP-Cy5.5 | MQ1-17H12 | BD |
| **IL-17** | PE | eBio64CAP17 | eBioscience |
| **IFNγ** | Alexa Fluor 700 | B27 | BD |
| **TNFα** | APC | Mab11 | BD |
| **Granzyme B** | FITC | GB11 | BD |

**Table S5.** Panel of antibodies used for AIM assay

| Marker | Fluorochrome | Clone | Manufacturer |
|---|---|---|---|
| **Viability** | Aqua | N/A | Invitrogen |
| **CD3** | BUV395 | SP34-2 | BD |
| **CD4** | FITC | L200 | BD |
| **CD8** | BV786 | RPA-T8 | BD |
| **CD20** | APC-H7 | 2H7 | BD |
| **CD134** | PE | L106 | BD |
| **CD137** | BV650 | 4B4-1 | BioLegend |
| **CD183** | BB700 | 1C6/CXCR6 | BD |
| **CD185** | PE-Cy7 | MU5UBEE | Invitrogen |
| **CD196** | PE-CF594 | 11A9 | BD |
| **CD279** | BV421 | EH12.2H7 | BD |
